# Multimodal profiling of lung granulomas reveals cellular correlates of tuberculosis control

**DOI:** 10.1101/2020.10.24.352492

**Authors:** Hannah P. Gideon, Travis K. Hughes, Constantine N. Tzouanas, Marc H. Wadsworth, Ang Andy Tu, Todd M. Gierahn, Joshua M. Peters, Forrest F. Hopkins, Jun-Rong Wei, Conner Kummerlowe, Nicole L. Grant, Kievershen Nargan, Jia Yao Phuah, H. Jacob Borish, Pauline Maiello, Alexander G. White, Caylin G. Winchell, Sarah K. Nyquist, Sharie Keanne C. Ganchua, Amy Myers, Kush V Patel, Cassaundra L. Ameel, Catherine T. Cochran, Samira Ibrahim, Jaime A Tomko, Lonnie James Frye, Jacob M. Rosenberg, Angela Shih, Michael Chao, Charles A. Scanga, Jose Ordovas-Montanes, Bonnie Berger, Joshua T. Mattila, Rajhmun Madansein, J. Christopher Love, Philana Ling Lin, Alasdair Leslie, Samuel M. Behar, Bryan Bryson, JoAnne L Flynn, Sarah M. Fortune, Alex K. Shalek

## Abstract

*Mycobacterium tuberculosis* lung infection results in a complex multicellular structure, the granuloma. In some granulomas, immune activity promotes bacterial clearance; in others, bacteria persist and grow. We identified correlates of bacterial control in cynomolgus macaque lung granulomas by co-registering longitudinal PET-CT imaging, single-cell RNA-sequencing, and measures of bacterial clearance. We find that bacterial persistence occurs in granulomas enriched for mast, endothelial, fibroblast and plasma cells, signaling amongst themselves via Type II immunity and wound healing pathways. In contrast, these interactions are largely absent in granulomas that drive bacterial control, which are often those that form later in the course of infection; these restrictive lesions are characterized by cellular ecosystems enriched for Type1-Type17, stem-like, and cytotoxic T cells engaged in pro-inflammatory signaling networks that involve diverse myeloid and non-immune cell populations. There is also a temporal aspect to bacterial control, in that granulomas that arise later in infection (in the context of an established immune response) share the functional characteristics of restrictive granulomas and are more capable of killing Mtb. Taken together, our results define the complex multicellular ecosystems underlying (lack of) granuloma resolution and highlight host immune targets that can be leveraged to develop new vaccine and therapeutic strategies for TB.

**One-Sentence Summary:** Bacterial control in TB lung granulomas correlates with distinct cellular immune microenvironments and time of formation after infection.

## Introduction

Tuberculosis (TB), caused by *Mycobacterium tuberculosis* (Mtb), remains a major global health threat (WHO, 2019). Mtb infection is characterized by the formation of granulomas predominantly in the lungs and lymph nodes (Flynn, 2010; Lin et al., 2014b; Russell et al., 2010; Ulrichs and Kaufmann, 2006). These spatially organized structures, composed of a mixture of immune and non-immune cells (Ehlers and Schaible, 2012; Flynn, 2010; Gideon et al., 2019; Lin et al., 2006; Mattila et al., 2013; Pagan and Ramakrishnan, 2014; Phuah et al., 2012; Reece and Kaufmann, 2012; Ulrichs and Kaufmann, 2006), are key sites of host-pathogen interactions which can either restrict or facilitate bacterial survival **(Fig S1A)**. Understanding the cellular and molecular features in granulomas that are associated with bacterial restriction versus failure to control infection is critical for the development of next-generation treatments and vaccines for TB. Delineating such protective responses in humans has been challenging given the limited accessibility of affected lung tissue and difficulty determining the true extent of bacterial control. The cynomolgus macaque model of Mtb infection, which recapitulates the diversity of human infection outcomes and granuloma pathologies, has been a transformative advance in the field, enabling detailed studies of the features of immunologic success and failure in Mtb granulomas (Canetti, 1955; Flynn, 2010; Lin *et al*., 2006).

A spectrum of granuloma types, organization and cellular composition has been described in both humans and non-human primates (NHP) (Canetti, 1955; Flynn, 2010; Hunter, 2011; 2016; Lin *et al*., 2006). Studies of Mtb infection in NHP have demonstrated that individual granulomas are dynamic (Coleman et al., 2014b; Lin et al., 2013; Lin *et al*., 2014b), changing in response to evolving interactions between bacteria and diverse host cell types (Ehlers and Schaible, 2012; Flynn, 2010; Flynn et al., 2003; Mattila *et al*., 2013; Phuah *et al*., 2012; Ulrichs and Kaufmann, 2006). The bacterial burden in individual granulomas is highest early in infection and then decreases due to increased bacterial killing as the immune response matures, even in animals that ultimately develop active TB (**Fig S1B-C)** (Cadena et al., 2016; Lin *et al*., 2014b; Maiello et al., 2018). Strikingly, however, individual granulomas within a single host follow independent trajectories with respect to inflammation, cellular composition, reactivation risk, and ability to kill Mtb (Coleman *et al*., 2014b; Gideon et al., 2015; Lenaerts et al., 2015; Lin *et al*., 2013; Lin *et al*., 2014b; Malherbe et al., 2016; Martin et al., 2017). We and others have profiled immune responses among individual cell types in macaque lung granulomas, including those of T cells (Diedrich et al., 2020; Foreman et al., 2016; Gideon *et al*., 2015; Lin et al., 2012; Mattila et al., 2011; Wong et al., 2018), macrophages (Mattila *et al*., 2013), B cells (Phuah et al., 2016; Phuah *et al*., 2012), and neutrophils (Gideon *et al*., 2019; Mattila et al., 2015), and also examined the instructive roles of cytokines, including IFN-γ, IL-2, TNF, IL-17 and IL-10 (Gideon *et al*., 2015; Lin et al., 2010; Wong et al., 2020). While these analyses have enabled key insights into how specific canonical cell types and effector molecules relate to bacterial burden, they have been relatively narrow and directed in focus, and have not revealed how the integrated actions of diverse cell types within individual granulomas influence control.

The emergence of high-throughput single-cell genomic profiling methods affords transformative opportunities to define the cell types, phenotypic states and intercellular circuits that comprise granulomas and inform their dynamics (Prakadan et al., 2017). Here, we developed and applied a multifactorial profiling pipeline—integrating longitudinal PET-CT imaging, single-cell RNA-sequencing (scRNA-seq)-based immunophenotyping, molecular measures of bacterial killing with immunohistochemistry and flow cytometry—to identify features of TB lung granulomas that correlate with bacterial clearance in cynomolgus macaques (**Fig 1A**). Leveraging it, we define the general cellular compositions and specific cell states associated with bacterial persistence or control. We further uncover TB-associated intercellular signaling networks and how they differ across granulomas that have different levels of bacterial clearance, identifying distinct participating cell types and pathways implicated in bacterial persistence or control. Collectively, our data define the cellular environments and holistic interaction networks within TB lung granulomas in which Mtb is controlled or alternatively survives and multiplies, nominating novel therapeutic and prophylactic targets for future investigation.

**Figure 1.**
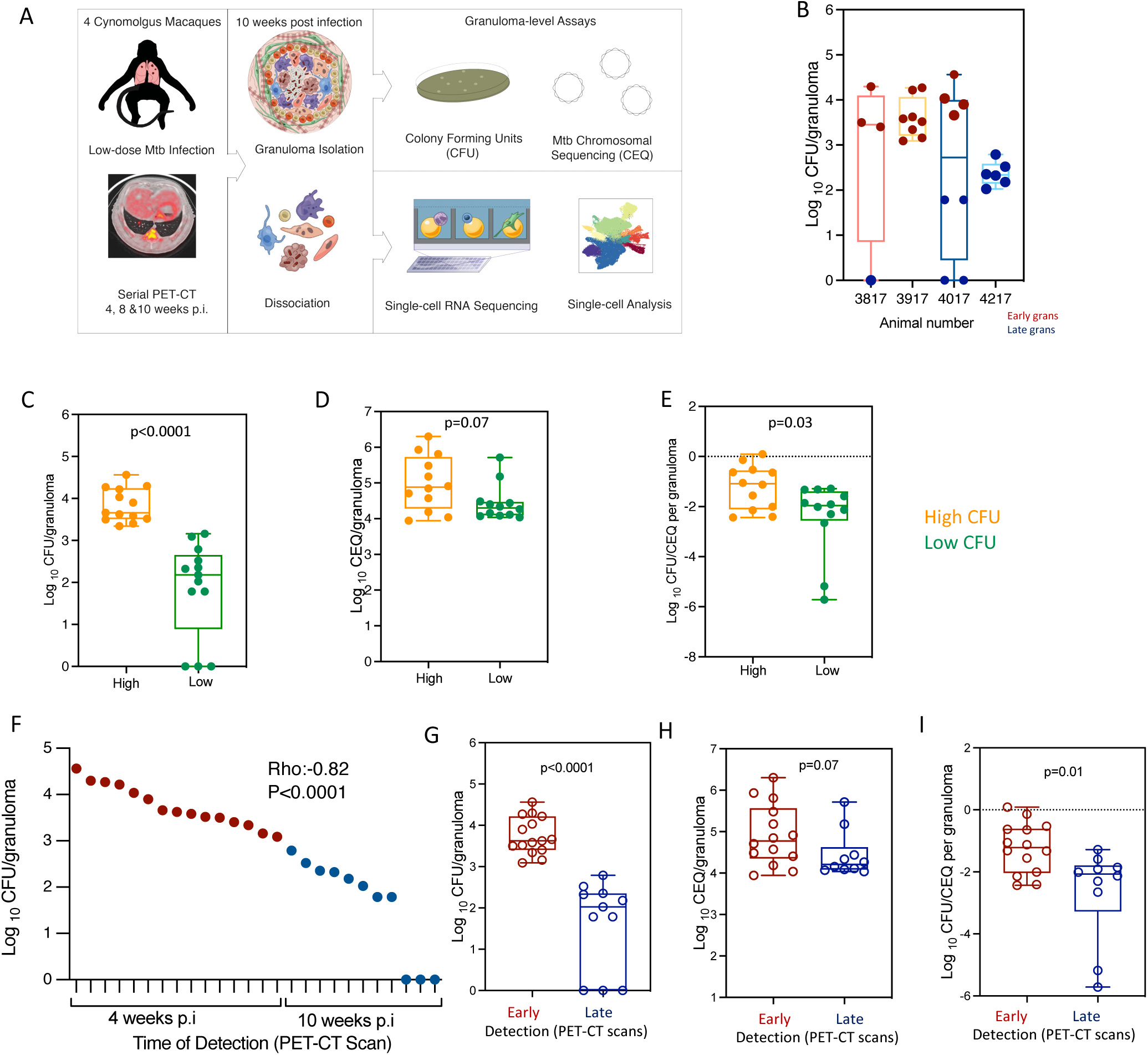
Study design, experimental set up, characteristics of animals over the course of Mtb infection and granuloma bacterial burden. (**A**) Study design: Cynomolgus macaques (n=4) were infected with a low-dose inoculum of Mtb (Erdman strain) and serial PET-CT scans were performed at 4, 8, and 10 weeks post-infection with the final scan used as a map for lesion identification at necropsy. Individual granulomas were excised and homogenized. CFU and CEQ assays were performed on all granulomas (top right) and 26 individual granulomas across 4 animals were randomly selected at necropsy for Seq-Well assays (bottom right). (**B**) Distribution of CFU per granuloma sampled for Seq-Well assay for each animal. Each dot is an individual granuloma. (**C**) /(**G**) CFU log10 per granuloma (total live bacteria); (**D**)/ (**H**) CEQ log10 per granuloma (Chromosomal equivalents, CEQ, live + dead Mtb) organized by time of detection; (**E**)/(**I**) Ratio between CFU (viable bacteria) and CEQ (total bacterial burden) i.e., relative bacterial survival. Lower ratio (negative values) corresponds to increased killing and higher ratio corresponds to increased Mtb survival. (**C-E**) organized by bacterial burden: low (Green); high (orange). (F) Individual Granuloma bacterial burden (log CFU) ploted with time of detection by PET-CT scans: 4 weeks post infection (early) or 10 weeks post infection (late). The granulomas in X axis is arranged in order of bacterial burden and time of detection. (**F-I**) time of detection by PET-CT scan (Table S1): early granulomas (maroon), late granulomas (blue). Each symbol is a granuloma. Box plot showing median, IQR and range. Mann Whitney U for panels **E-G**.

## Results

We sought to define the complex cellular ecosystems of granulomas that manifest different degrees of bacterial control in NHP. Four cynomolgus macaques were infected with a low dose of Mtb (<10 CFU; Erdman strain) and followed for 10 weeks (**Fig 1A**). 10 weeks post-infection was chosen as a pivotal time point in which bacterial killing can be identified in some but not all granulomas (**Fig S1B**), providing the potential to examine a range of bacterial burdens across granulomas in our analyses. Progression of Mtb infection and individual granuloma dynamics were monitored at 4, 8, and 10 weeks post infection (p.i.) using PET-CT imaging of FDG avidity as a proxy for inflammation **(Fig S1D-E, Table S1)** (Coleman *et al*., 2014b; White et al., 2017). At necropsy, individual PET-CT identified lung granulomas were excised and dissociated to obtain a single-cell suspension; viable bacterial burden (CFU, colony forming units – i.e., culturable live bacterial burden) and cumulative (live + dead) bacterial load (chromosomal equivalents, CEQ) were measured to define the extent of bacterial growth and killing in each granuloma (Lin *et al*., 2014b; Munoz-Elias et al., 2005).

Twenty-six granulomas from these four animals were randomly selected at the time of necropsy for scRNA-seq analysis. Among the 26, there was a range of granuloma-level bacterial burdens, from sterile (0 CFU/granuloma) to high (4.6 log_10_ CFU/granuloma) (**Fig 1B-C**; **Table S1**). The granulomas were binned based on bacterial burden (low, n=13 and high, n=13). There was a significant difference in CFU between low and high CFU granulomas (median 2.2 (low) vs 3.6 (high) log_10_ CFU/granuloma, p<0.0001, Mann Whitney U test) (**Fig 1C**). To determine whether low CFU reflected reduced bacterial growth or increased bacterial killing, we assessed the total number of bacterial genomes (CEQ), where we have previously shown that the genomes of dead bacteria are not readily cleared and that CEQ provides a measure of cumulative bacterial load (Munoz-Elias *et al*., 2005). We observed no significant difference in CEQ values between low and high burden granulomas, indicating that the granulomas supported roughly similar cumulative Mtb growth over the course of infection (**Fig 1D**). However, the extent of bacterial killing, calculated as the ratio of CFU to CEQ, was significantly higher in the low bacterial burden granulomas (**Fig 1E**), indicating that the lower CFU reflected greater killing rather than more limited bacterial growth.

We then sought to identify granuloma features correlated with the degree of bacterial control. *Post-hoc* analysis of serial PET CT imaging data revealed a strong association between the apparent timing of lesion formation and the extent of bacterial control. All the high bacterial burden granulomas were detected at the 4-week scan, while most (11/13) of the low bacterial burden granulomas were first detected at the final pre-necropsy scan (10 weeks) (**Fig S1E,** **Fig 1F-G**). Consistent with these data, we further evaluated bacterial burden between early and late appearing granulomas in a total of 10 animals at 10 weeks p.i. (**Fig S1G-H**) and again found that the median CFU/granuloma per animal was significantly lower in late granulomas as compared to early ones. We considered the model that late lesions have lower CFU because the bacterial population has simply not had sufficient time to expand. However, since the cumulative bacterial burdens (CEQ) in early and late lesions were not significantly different (**Fig 1H**), the data are consistent with more bacterial killing in late appearing granulomas (−2.1 log_10_ CFU/CEQ per granuloma) as compared to early appearing ones (−1.2 log_10_ CFU/CEQ per granuloma, p=0.01, Mann Whitney U test) (**Figure 1I**). Late appearing granulomas could be due to differences in the timing of lesion formation, most likely due to a dissemination event from an early granuloma, such that bacterial replication occurred in the context of an activated immune response (Martin *et al*., 2017) or differences in the characteristics of the initial inflammatory response such that late appearing granulomas were not detectable by imaging until later in infection.

### Cellular composition of TB lung granulomas

To identify cellular and molecular factors associated with increased Mtb killing in an unbiased fashion, we loaded a single-cell suspension from each of the 26 granulomas onto a Seq-Well array (Gierahn et al., 2017) under Biosafety Level 3 conditions, and then processed and sequenced as previously described (Gierahn *et al*., 2017). After aligning the data to the *Macaca fascicularis* (cynomolgus macaque) genome and performing robust quality controls and granuloma-specific technical corrections, we retained 109,584 high-quality single-cell transcriptomes for downstream analysis (**Fig S2; Table S2**).

Among these, we resolved 13 general cell types (**Fig 2A,B** and **Fig S3A-G**) through dimensionality reduction, Louvain clustering, and examination of canonical lineage defining genes and reference signatures from the Tabula Muris (Tabula Muris et al., 2018), Mouse Cell Atlas (Han et al., 2018) and SaVanT database (Lopez et al., 2017) (**Fig S3 A-G, Table S3**). These 13 encompass groups of lymphocytes, including B cells (defined by expression of *MS4A1*, *CD79B*, & *BANK1*), T and NK cells (T/NK; *GNLY, TRAC, CD3D,* & *GZMH*) and plasma cells (*IGHG1* & *JCHAIN)*); myeloid cells, including conventional dendritic cells (cDCs; *CLEC9A, CST3,* & *CPVL*), plasmacytoid dendritic cells (pDCs; *LILRA4* and *IRF8*), and macrophages (*APOC1, LYZ,* and *APOE*); mast cells (*CPA3* & *TPSAB1*); neutrophils (*CCL2, CXCL8,* & *CSF3R*); erythroid cells (*HBA1* & *HBB*); stromal cells, including endothelial cells (*RNASE1, EPAS1,* & *FCN3*) and fibroblasts (*COL3A1, COL1A1,* & *DCN)*; Type-1 pneumocytes (*AGER*); and, Type-2 pneumocytes (*SFTPC, SFTPB,* and *SFTPA1*) (**Fig 2A** **& B, Fig S3G** and **Table S3 & S4**). For each of the 13 cell types, we also performed further within cell-type sub-clustering; in these analyses, we only detected substructure among the T/NK and macrophage clusters (detailed below, **Methods**).

**Figure 2.**
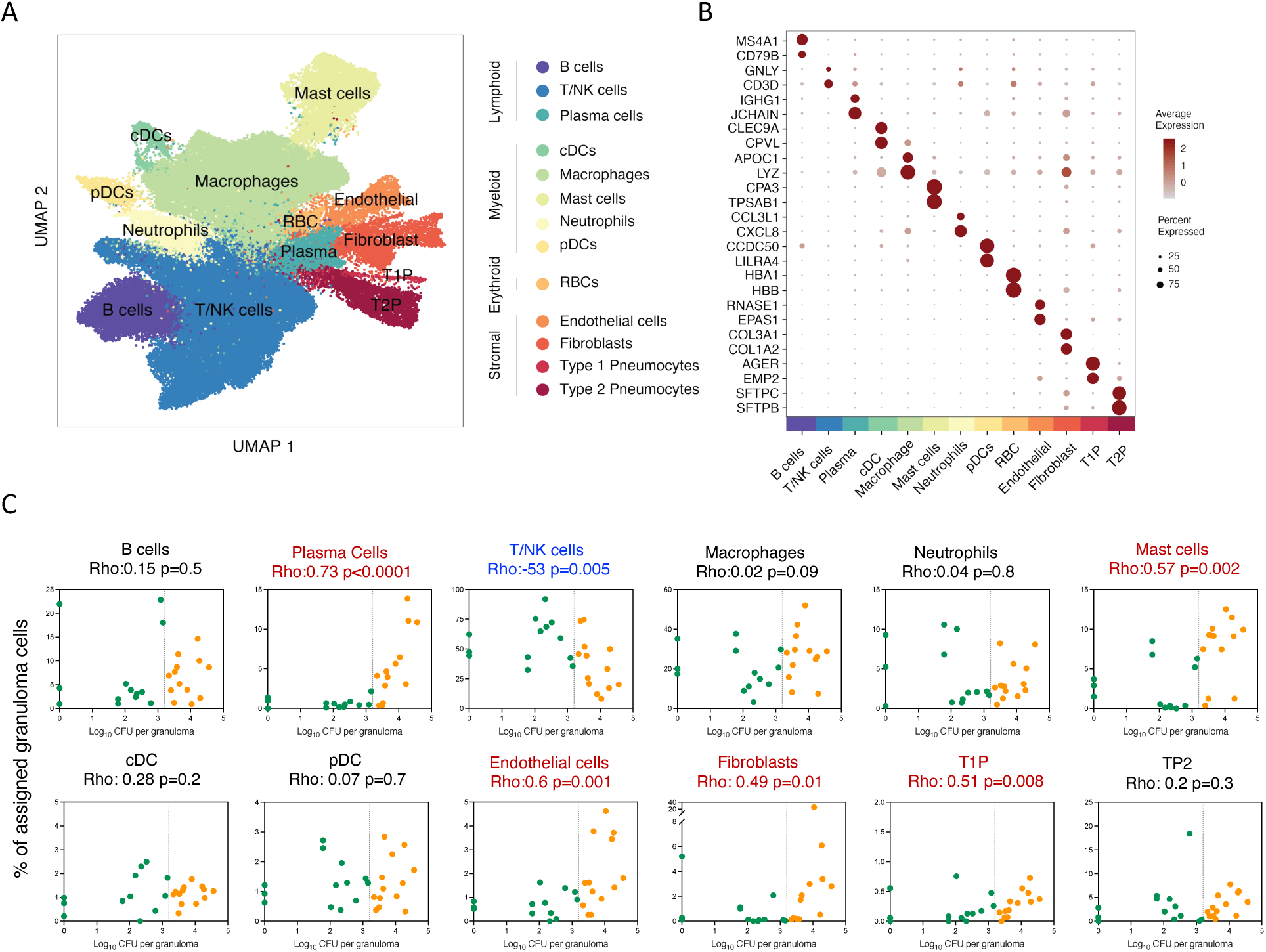
Analysis of single-cell sequencing of tuberculosis lung granulomas. (**A**) UMAP plot of 109,584 cells from 26 granulomas colored by identities of 13 generic cell types. **(B)** Expression levels of cluster defining genes enriched across 13 generic cell types. Color intensity corresponds to the level of gene expression, while the size of dots represents the percent of cells with non-zero expression in each cluster. **(C)** Significant correlations between proportion of canonical cell types with bacterial burden of individual granulomas (Log10 CFU per granuloma) using non-parametric Spearman’s rho correlation test. Color indicated binned granuloma bacterial burden: low (green) and high (orange).

### Cell types associated with timing of granuloma formation and control

To investigate the relationship between cell type composition and bacterial burden, we quantified the correlation between cellular frequency and CFU across all granulomas. We considered cellular frequencies in granulomas as a function of viable bacterial burden. Our data reveal multiple cell types that are significantly enriched in higher bacterial burden (early appearing) granulomas, including plasma cells (relative cell abundance vs CFU, p<0.0001, non-parametric Spearman’s rho correlation test), mast cells (p=0.002), endothelial cells (p=0.001) and fibroblasts (p=0.011) (**Fig 2C, Table S5**). By contrast, T/NK cells were more abundant in lower bacterial burden (late appearing) granulomas (p=0.0055) (**Fig 2C, Table S5**). Cynomolgus macaques are variable in their infection outcomes (**Fig 1B**), so to control for inter-subject variability, each of the cellular associations between granuloma dynamics and bacterial control was examined both across all animals and lesions, and through a directed analysis of the granulomas from a single NHP host (4017) (**Fig S3H**). We further confirmed these trends by performing deconvolution on bulk RNA-sequencing data of 12 additional granulomas (6 high CFU (early) and 6 low (late) bacterial burden granulomas) from separate macaques (**Fig S4A**).

### High bacterial burden granulomas are characterized by fibrosis and Type II immune features

The presence and function of mast cells in Mtb lung granulomas has not been previously described. Therefore, to validate this observation, we performed immunohistochemistry on NHP and human granuloma sections using Tryptase and C-kit/CD117 markers (**Fig S4D & E**). This confirmed the presence of mast cells within both NHP and human granulomas, and further revealed that they primarily localize to the outer regions of NHP granulomas, including the lymphocyte cuff (**Fig S4D**), and can be found within and around human granulomas (**Fig S4E**). In our data, mast cells are distinguished by their expression of *IL4* and *IL13* (**Fig S4B**), which we also recently observed in a study of human nasal polyposis, a Type II inflammatory disease associated with dramatic epithelial remodeling (Ordovas-Montanes et al., 2018). Mast cells are also marked by expression of *ALOX5A* and *ALOX5AP*, which encode the system to synthesize the anti-inflammatory lipoxin LXA4; the balance between LXA4 and the pro-inflammatory lipoxin LTB_4_ has been strongly implicated in the progression of TB disease in humans (Tobin et al., 2012; Tobin et al., 2010).

Plasma cells are also abundant in high burden lesions, consistent with previous findings (Jacobs et al., 2016; Phuah *et al*., 2012). Recruitment of mast cells can be characteristic of allergic Type II immune responses mediated by IgE (Kanagaratham et al., 2020), but mast cell function is also regulated by IgG, which is much more abundant in the circulation and tissues. Among the plasma cells in our scRNA-seq dataset, the vast majority express either *IGHG* or *IGHA* (Collins and Jackson, 2013) constant chains (**Fig S4B, C)**, suggesting that IgG and IgA are the dominant antibody classes induced by Mtb infection in the granuloma microenvironment. Taken together, these data suggest that granulomas with failed bacterial clearance are characterized by a Type II immune environment, but the antibody features are not consistent with a canonical allergic response.

### T and NK functional subclusters as mediators of protection

Of the 13 broad cell types, only the T/NK cell subcluster is associated with more robust bacterial control in granulomas (p=0.0055, non-parametric Spearman’s rho correlation test) (**Fig 2C**). To further assess functional diversity within the 41,622 cells that comprise the T and NK cell cluster and their association with bacterial burden, we performed additional sub-clustering analyses. This revealed 13 T/NK cell subclusters which we annotated based upon expression of lineage defining markers, known cytotoxic, regulatory and proliferation genes (**Fig 3A, C** and **S5, Tables 1 and S6)** and TCR constant gene (*TRAC*, *TRBC*, and *TRDC*) expression (**Fig 3B**). The process of annotation revealed that most subclusters did not correspond neatly to canonical T and NK cell subsets, consistent with recent studies in other systems (Rath et al., 2020). Where possible, we annotated each based on known T cell markers and literature-derived genes of interest; we note that these genes are parts of broader transcriptional signatures that appear to reflect dominant cellular response states superimposed on cell lineage-associated gene expression programs. Among the 13 T/NK cell subclusters, 6 were significantly negatively associated with bacterial burden (**Fig 3D, Table S5**).

**Figure 3.**
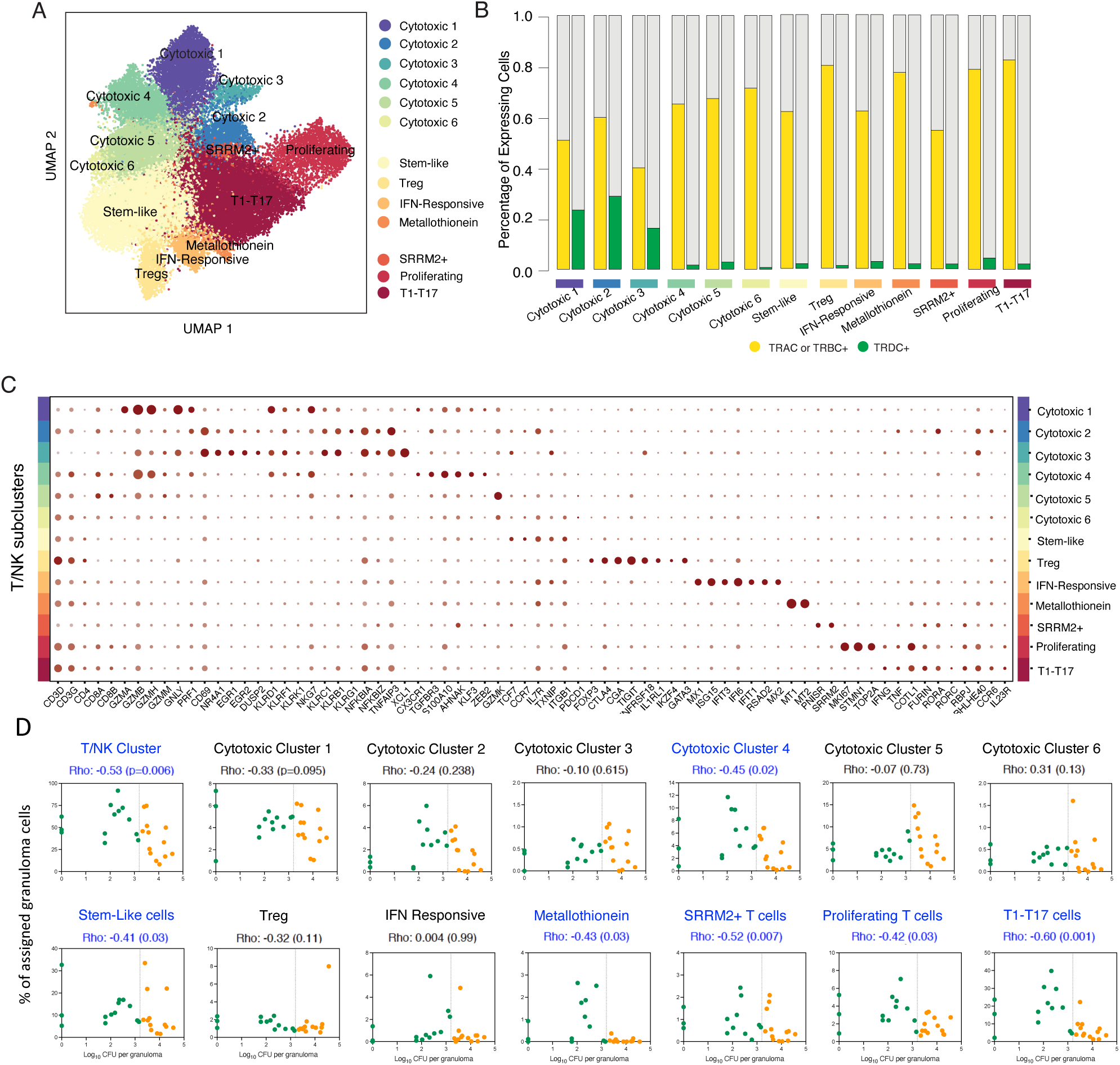
Diversity in the unified T and NK cell cluster and relationship to granuloma-level bacterial burden. **(A)** Subclustering 41,222 cells in the unified T/NK cell cluster, colored by subclusters. Subclusters are numbered based the expression patterns. **(B)** Frequency of expression of TCR genes *TRAC*, *TRBC1* or *TRBC2* (yellow) and *TRDC* (green) across 13 T/NK cell subclusters. **(C)** Expression levels of T/NK cell cluster-defining genes. Color intensity corresponds to the level of gene expression and the size of dots represents the percent of cells with non-zero expression in each cluster. Y-axis identifies subclusters. **(D)** Significant correlations between proportion of T/NK subclusters with bacterial burden of individual granulomas (Log10 CFU per granuloma) using non-parametric Spearman’s rho correlation test. Color indicated binned granuloma bacterial burden: low (green) and high (orange).

### A prominent role for Type1-Type 17 T cells in bacterial control

One T/NK cell subcluster represented the most abundant cell type identified across all granulomas (8.8%) (**Table S4**) and the strongest correlate with bacterial control (p=0.001, non-parametric Spearman’s rho correlation test) (**Fig 3D**; **Table S4 & S5**). This subcluster, which we designated Type1-Type17 (T1-T17) (**Fig 3C**), is enriched for expression of classical Th1-associated genes, including *IFNG* and *TNF* (Raphael et al., 2015), as well as transcription factors associated with Th17 differentiation (Yosef et al., 2013), including *RORA* (Yang et al., 2008)*, RORC* (Ivanov et al., 2006), *RBPJ* (Meyer Zu Horste et al., 2016), and *BHLHE40* (Huynh et al., 2018; Lin et al., 2016; Lin et al., 2014a). While we also detected additional features of T17 cells, including *CCR6* (Hirota et al., 2007) and *IL23R (Kobayashi et al., 2008)*, we did not observe expression of either *IL17A* or *IL17F* (**Fig 4A; Table S6-7)**. Collectively, this hybrid gene expression state is consistent with previously described expression programs for Th1* or ex-Th17 cells, which are believed to be precursors to tissue resident memory cells (Amezcua Vesely et al., 2019). Previous studies have revealed a prominent role for CD4 Th1 and Th17 cytokines in control of Mtb infection, including IFN-γ, TNF, and IL-17 (Algood et al., 2005; Green et al., 2013; Khader et al., 2007; Khader and Gopal, 2010; Lin et al., 2007; Lyadova and Panteleev, 2015; Millington et al., 2007; O’Garra et al., 2013; Scriba et al., 2017), and studies in NHP granulomas suggest an association between T1 and T17 cytokine expression and bacterial burden (Gideon *et al*., 2015). In addition, in murine models, BHLHE40 is required for control of Mtb infection, as a repressor of IL-10 production (Huynh *et al*., 2018). Notably, while Th1* and ex-Th17 subsets are described primarily as CD4 T cells (Darrah et al., 2020; Gideon *et al*., 2015; Lyadova and Panteleev, 2015; Mpande et al., 2018), our T1-T17 sub-cluster is characterized by the expression of both *CD4* and *CD8A/B* transcripts (**Fig 3C and 4C, Fig S5D-E)**.

**Figure 4.**
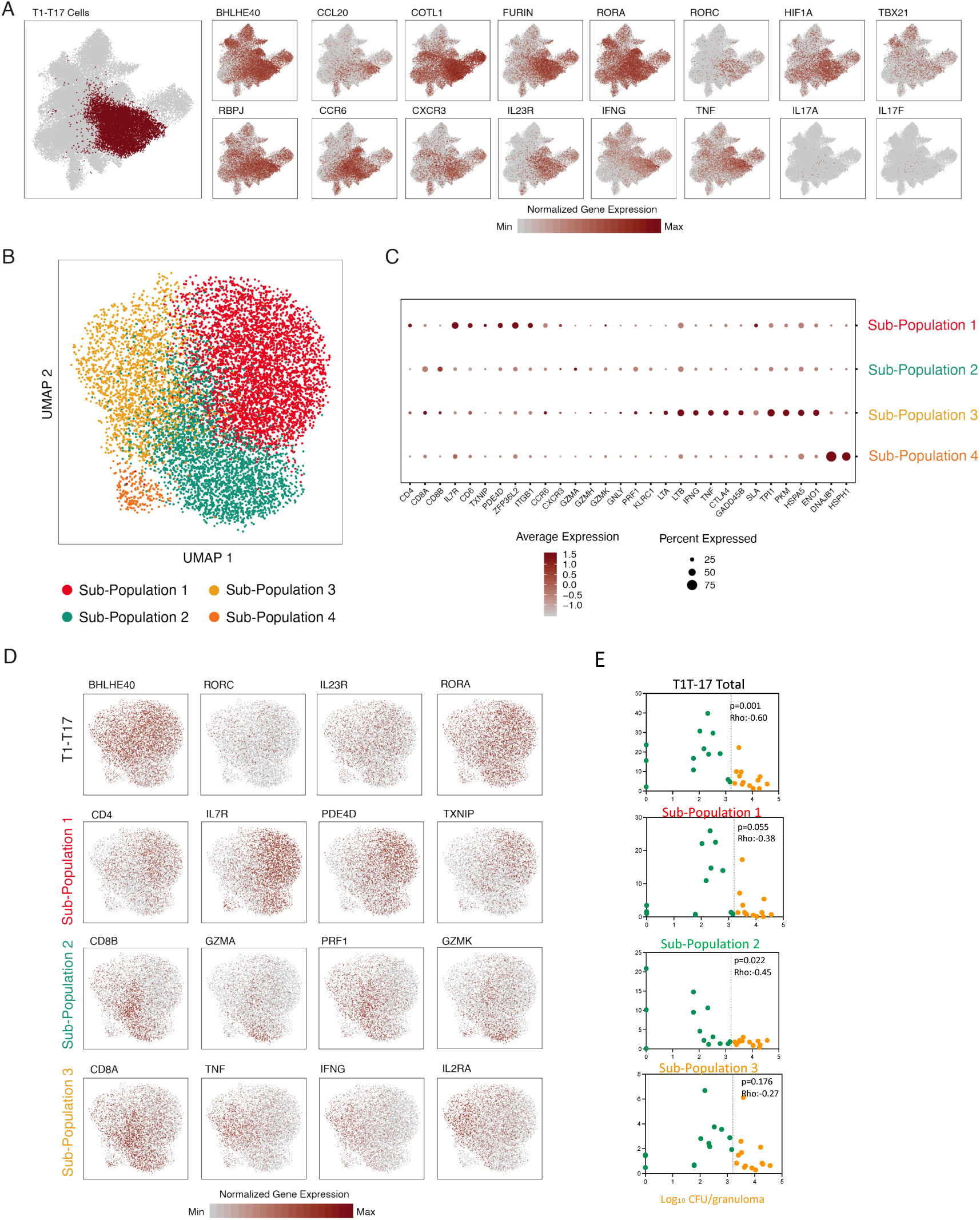
Phenotypic Diversity in T1-T17 cells. **(A)** T1-T17 subcluster overlaid on unified T/NK cell cluster (left) and colored by normalized expression values for T1-T17 subcluster-defining genes (bold outlined boxes) and non-enriched canonical Type1 and Type 17 genes (right). **(B)** Subclustering of 9,234 T1-T17 cells resulting in 4 phenotypic sub-populations. **(C)** Cluster defining genes for T1-T17 subpopulation 1, 2, 3 and 4. Color intensity corresponds to the level of gene expression and the size of dots represents the percent of cells with non-zero expression in each cluster. **(D)** Subclustering of T1-T17 cells colored by normalized gene expression values for selected subcluster (top row) and sub-population defining genes. **(E)** Significant correlations between proportion of T1-T17 subcluster and subpopulations with bacterial burden of individual granulomas (Log10 CFU per granuloma) using non-parametric Spearman’s rho correlation test.

To better resolve the identities of the cells in this cluster, we further sub-clustered the 9,234 T1-T17 cells. This revealed 4 distinct subpopulations, each of which expressed T1-T17 cluster markers (*RORA, RORC, IL23R*, and *BHLHE40*) but were further distinguished by markers of cell type and state **(Fig 4B, Table S7**): T1-T17 subpopulation 1 is distinguished by expression of *CD4* and markers of activation and motility, including *IL7R*, *CD6*, *TXNIP*, *PDE4D*, *ZFP36L2*, *ITGB1*, *CCR6*, and *CXCR3* (**Fig 4B,C; Tables 1 and S7**), making it most akin to ex-Th17 cells; T1-T17 subpopulation 2 is characterized by increased relative expression of both *CD8A* and *CD8B* and cytotoxic effector molecules; T1-T17 subpopulation 3, which includes cells expressing either *CD8A/B* or *CD4*, is characterized by cytokine gene expression (*IFNG, TNF, LTA*, and *LTB*) and markers of an inhibitory cell state (*CTLA4*, *GADD45B*, and *SLA*); T1-T17 subpopulation 4 is very low in abundance and characterized by heat shock and DNA damage associated transcripts (*DNAJB1* and *HSPH1*). There was a trend towards negative association between bacterial burden and higher abundance of T1-T17 subpopulation 1 (p=0.055, non-parametric Spearman’s rho correlation test) and a significant negative association between bacterial burden and abundance of T1-T17 subpopulation 2 (p=0.02). Surprisingly, T1-T17 subpopulation 3 was not correlated with bacterial burden, despite expressing elevated levels of *IFNG* and *TNF* (**Fig 4E**, **Table S5**), cytokines generally considered as critical mediators of control in Mtb infection (O’Garra *et al*., 2013; Scriba *et al*., 2017).

### CD4 and CD8 subclusters associated with low bacterial burden

Among the remaining 12 T/NK cell subclusters, 6 are enriched for both CD4 and CD8 expression (**Fig 3A-C**, **Fig S5D&E**, **Table 1, S6**). Of these, 5 are significantly associated with more robust bacterial control (**Figure 3D** **& S5D-E**). We annotated the most abundant of these as stem-like T cells (8.3% of granuloma cells, p=0.03 non-parametric Spearman’s rho correlation test, **Fig 3D, Table S5)** based on elevated expression of markers of naïve and memory T cells (*TCF7*, *CCR7, IL7R,* and *TXNIP*) and activation or memory state (*CD69* and *ITGB1*) (**Fig 3C, Table S6**). These cells may represent a “stem-like” population of T cells, which has been described as an early differentiating memory phenotype, distinct from naïve T cells, that are long-lived and possess a unique ability to proliferate and self-renew (Ahmed et al., 2016; Caccamo et al., 2018; Gattinoni et al., 2011). The second CD4/CD8 subcluster associated with control contains proliferating T cells (2.4%; p=0.03; **Fig 3D, Table S5**) and is characterized by high expression of transcripts associated with cellular proliferation (*MKI67*, *STMN1*, and *TOP2A*) **(Fig 3C, Table S6)**, consistent with published data that T cell proliferation occurs within NHP and human granulomas (Gideon *et al*., 2015; McCaffrey et al., 2020; Ohtani, 2013; Phuah *et al*., 2016; Phuah *et al*., 2012; Wong *et al*., 2018). The third is a very small population of Metallothionein expressing T cells (0.05%; p=0.03; **Fig 3D, Table S5**), defined by metallothionein genes, such as *MT1* and *MT2* **(Fig 3C, Table S6),** which play a role in negative regulation of Type 1 regulatory (Tr1) CD4+ cells (Wu et al., 2013). The fourth, SRRM2-T cells (0.6%, p=0.007), is characterized by enrichment of genes associated with nuclear speckles and splicing factors such as *PNISR* and *SRRM2* **(Figure 3C&D, Table S5-6)**, the latter of which has been associated with alternate splicing in Parkinson disease (Shehadeh et al., 2010) and has a critical role in the structural organization of the genome (Hu et al., 2019).

The remaining two CD4/CD8 subclusters are not associated with bacterial control. Interestingly, one is regulatory T cells (1.2%), defined by elevated expression of canonical Treg markers (*FOXP3, CTLA4, TIGIT,* and *IL1RL1*) and *GATA3*, a Th2 lineage-defining transcription factor that has been observed in a subset of tissue-resident Tregs (Fig 3C&D**, Table S5-6)**. The final subcluster is interferon responsive T cells (0.4%), which are enriched for Type-I interferon inducible molecules (*MX1*, *ISG15*, *IFIT3*, *IFI6*, *IFIT1*, *RSAD2*, and *MX2*) *(Szabo et al., 2019)* **(Fig 3C-D, Table S5-6).**

### Bacterial control is associated with a specific cytotoxic T cell population

The remaining 6 T/NK subclusters are broadly defined by expression of *CD8A* and/or *CD8B* and cytotoxic genes, including granzymes (*GZMA, GZMB, GZMH, GZMK*, and *GZMM*), granulysin (*GNLY)*, and/or perforin (*PRF1*) (designated Cytotoxic 1-6, **Fig 3C, Table 1**). We confirmed expression of multiple granzymes among CD8αβ T cells in Mtb granulomas by flow cytometry (**Fig S6**) from animals in other ongoing studies.

Low bacterial burden granulomas are associated with a higher proportion of cells from cytotoxic subcluster C4 (3.8% of granuloma cells; p=0.02, non-parametric Spearman’s rho correlation test) (**Fig 3D; Table S5**). C4 expresses both *CD8A* and *CD8B* and *TCRA* and *TCRB*, but not *TCRD,* indicating that it is composed primarily of conventional CD8αβ T cells (**Fig 3B&C, S5D)**. C4 is further enriched for genes associated with cytotoxic effector functions (*PRF1, GZMH*, *GZMB,* and *GZMM)*, motility, migration and tissue residency (*CX3CR1, TGFBR3*, and *S100A10*), and regulators of cell state (*AHNAK*, *KLF3*, and *ZEB2*; **Fig 3C, Table S6**).

The remaining 5 cytotoxic subclusters did not associate with bacterial control. Cytotoxic subclusters C1-3 are enriched for the expression of *CD8A* but not *CD8B* and elevated *TCRD*, implying that these cells possess innate cytotoxic function (**Fig 3B-C**). C1 is further characterized by high expression of cytotoxic effector genes*—GNLY* and *PRF1; GZMH*, *GZMA* and *GZMB;* as well as *KLRD1, KLRC1*, *KLRC2*, and *NKG7—*which suggests that subcluster 1 contains a greater proportion of highly cytotoxic innate CD8+ T cells (possibly NKT cells), γδ T cells, and NK cells (**Fig 3B-C, Table 1, S6**). C2 is also enriched for NK receptors and CD8 T cell activation markers in addition to a trio of transcription factors (*EGR1, EGR2*, and *DUSP2*) described to distinguish peripheral tolerant CD8 T cells (Schietinger et al., 2012) **(Fig 3B-C, Table 1, S6)**. C3 appears to be more selectively enriched for NK cells with elevated expression of cytotoxic and NK cell markers and low expression of *CD3D* and *CD3G.* C5, which like C4 expresses both *CD8A* and *CD8B* and *TCRA* and *TCRB*, but not *TCRD*, is distinguished by elevated expression of *GZMK* (**Fig 3C**); granzyme K expressing CD8 cells have been recently described as a hallmark of immune dysfunction in inflammation (Mogilenko et al., 2021). C6 was not detected in sufficient frequency (<0.3%) to draw meaningful conclusions. The functional complexity of these 6 subclusters, along with the common and distinct responses they represent, suggests a significant and underappreciated role for cytotoxic cells in TB granulomas.

### Macrophage heterogeneity in Mtb granulomas

While macrophages are responsible for much of the bacterial killing within granulomas, we did not observe any association between overall macrophage abundance and bacterial burden (**Fig 2** **and S7**). Yet, like the T/NK cell cluster, the macrophage cluster had discernable substructure based on unbiased gene expression analyses. Among the 27,670 macrophages, we identified 9 subclusters (**Table S8**), none of which were independently associated with bacterial control with the exception of Mac 4 (0.07%), a very small subpopulation of macrophages expressing *INSIG1* and *EREG* (p<0.0001) (**Fig S7E, Table S8**)).

### Cellular ecology of pulmonary TB granulomas

Given demonstrable differences in cellular composition across the bacterial burden spectrum, we wondered whether specific cell types significantly co-occur in TB lung granulomas and collectively influence control. We calculated the pairwise Pearson correlation matrix between all major cell types, subclusters, and subpopulations across the 26 granulomas (**Fig 5A**). Using hierarchical clustering of this pairwise correlation matrix, we defined 5 groups of cell types whose collective abundances are associated across granulomas (**Fig 5A**, **Table S9**). Of these, Group 2 (shown in red), which includes mast cells, plasma cells, macrophage subcluster 4 and certain stromal populations, is significantly expanded in high bacterial burden granulomas (Mann-Whitney U Test, p=3*10^-4^; **Fig 5B, Table S10, S11**). Group 3 (shown in blue) is significantly more abundant in low bacterial burden granulomas (p=0.026; **Fig 5B**, **Table S10, S11**) and consists of many T cell subclusters/subpopulations, including Stem-like, Cytotoxic subclusters C2, C4, & C6, Metallothionein, Proliferating, SRRM2+, T1-T17 subpopulations 1,3 and 4, as well as a single macrophage subset, Mac7. This macrophage subset is distinguished in part, by expression of the immunomodulatory genes *IDO* and *CHIT* (encoding chitotriosidase), which is abundantly produced by lipid-laden macrophages in other conditions such as Gaucher’s disease, Niemenn-Pick disease, and atherosclerosis (Barone et al., 2007; Yap et al., 2020).

**Figure 5.**
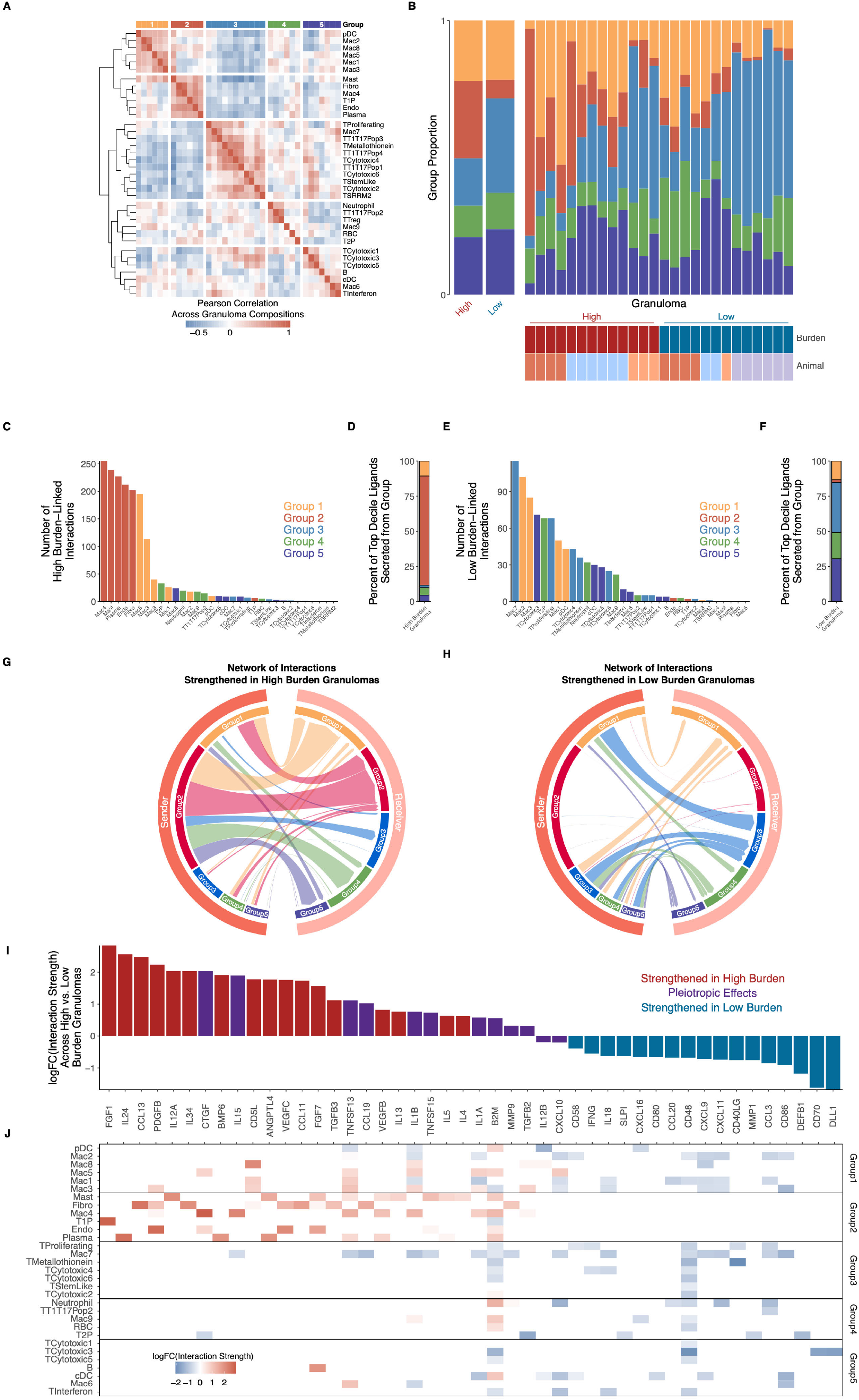
Cellular ecosystem in TB lung granulomas. **(A)** Pairwise Pearson correlation values proportions of canonical cell types and T/NK and macrophage subclusters across 26 granulomas. Hierarchical clustering of correlation coefficients identified 5 groups (indicated by color and number) of cell types with correlated abundance in granulomas. **(B)** Composition of each granuloma by cell type group. Left bar graph shows all high burden and all low burden granulomas grouped together, with right bar graph split by individual granuloma. **(C)** Number of interactions strengthened in high burden granulomas, organized by sender cell clusters (i.e., cell cluster producing the ligand). **(D)** Representation of each cell type group as sender cell population among the 10% of ligands most strengthened in high burden granulomas. **(E)** Number of interactions strengthened in low burden granulomas, organized by sender cell clusters. **(F)** Representation of each cell type group as sender cell population among the 10% of ligands most strengthened in low burden granulomas. **(G)** Network of interactions across cell type groups, subsetted to only highlight interactions strengthened in high burden granulomas. Widths of arcs are proportional to number of interactions between cell type groups, and widths are on same scale as for subfigure F. n = 2,586 statistically significant interactions, 1,715 of which were strengthened in high burden granulomas. **(H)** Network of interactions across cell type groups, subsetted to only highlight interactions strengthened in low burden granulomas. Widths of arcs are proportional to number of interactions between cell type groups, and widths are on same scale as for subfigure E. n = 2,586 statistically significant interactions, 871 of which were strengthened in high burden granulomas. **(I)** Overall high-vs-low granuloma burden fold-change of interactions strengths of key ligands, averaged across all statistically significant interactions. **(J)** Cell cluster-specific interaction strength fold changes of each ligand, averaged across all statistically significant interactions where each cell cluster was the sender population.

### Distinct cellular ecosystems associate with granuloma-level bacterial burden

To further explore how specific cellular compositions might constitute distinct tissue niches that support different levels of bacterial control, we examined putative cell-cell interactions within each granuloma. For each potential interacting cell-type pair, we constructed edge weights for receptor-ligand combinations, adjusting to account for differences in the abundance of the sender cell type, relative ligand/receptor expression, and the percent of receptor positive cells (**Methods**).

To obtain an initial view of cell-cell signaling across granulomas of different burden, we examined the extent and strengths of interactions across cell type groups. High bacterial burden lesions are dominated by signals sent by Group 2 cell types (i.e., mast, fibroblast, endothelial, plasma, type I pneumocyte, and macrophage subset 4); these cell types display the highest counts of high burden-linked interactions as well as those most strengthened in high burden granulomas (p < 2.2E-16, binomial test against null of all Groups having equal interaction likelihoods) (**Fig 5C-D**). In contrast, interactions in low burden granulomas more evenly involve Groups 1, 3, 4 and 5, with Group 3 showing the strongest enrichment for signaling activity strengthened in low burden granulomas (p = 1.2E-4, binomial test against null of all Groups having equal interaction likelihoods; p = 0.008, binomial test against null of equal interaction likelihoods among non-Group 2 cells) (**Fig 5E-F**). These contrasting patterns of intercellular communication suggest distinct signaling architectures underlying different degree of bacterial control, with Group 2 cells dominating activity within high-burden lesions, compared to coordinated signaling across Groups in low-burden cases.

We further examined shifts in intercellular interaction network topology by more comprehensively quantifying the sender and receiver activity associated with different levels of bacterial burden. This more directed investigation revealed significantly different patterns of intercellular signaling between high and low burden granulomas (p < 2.2E-16, Pearson’s chi-squared test). Subsetting all interactions to those strengthened in high burden granulomas, we find that Group 2 cell types are the key source of intercellular signals (i.e., senders in 67% of interactions strengthened in high burden granulomas) (**Fig 5G**). High burden lesions also exhibit strong intra-Group 2 signaling, with 58% of signals received by Group 2 cell types originating from Group 2 cell types themselves. This suggests that high burden lesions are driven by self-reinforcing interactions amongst Group 2 cell types (e.g., between mast cells, plasma cells, fibroblasts, and endothelial cells). In contrast, when subsetting to interactions strengthened in low burden granulomas, we find only sparse contributions from Group 2 cell types (**Fig 5H**); instead, low burden granulomas are characterized by a more even distribution of signals stemming and terminating in Group 1, 3, 4, and 5 cell types, suggestive of a coordinated immune response involving multiple cellular subsets (e.g., the T cell and macrophage subsets present in these Groups).

We next examined which specific axes of intercellular communication, and among whom, associate with varying levels of bacterial control. In looking more directly at the signals underlying the interaction networks that associate with burden, we find dramatic differences in intercellular crosstalk involving both canonical and non-canonical immune mediators that may impair or facilitate bacterial control. Among the ligands whose interactions are most strengthened in high burden granulomas, we identify genes implicated in fibrosis (e.g, *FGF1, PDGFB, CTGF, FGF7, IL34*), vascular remodeling (*VEGFB, VEGFC*, *ANGPTL4*) and TGFβ signaling (*TGFB3, BMP6*), suggestive of a wound healing response (**Fig 5I**) (Joshi et al., 2020; Padela et al., 2008). In addition, we observe evidence of intercellular communication via genes implicated in Type II immunity (*CCL11, CCL13, CD5L, IL4, IL5, IL13, IL24*) and allergy-linked inflammation (*CCL19*) (Nakano et al., 2019). We note that these specific ligands are largely produced and received by Group 2 cell types (with only sparse contributions from Groups 3-5). Collectively, this supports a model where intra-Group 2 signaling drives a self-reinforcing high burden microenvironment via wound healing-like responses and associated Type II immune activity (**Fig 5J**). This interpretation is further supported by an enrichment of pathways such as FGF, VEGFR, and PI3K signaling, as well as organogenesis and tissue remodeling processes (**Fig S8A**).

In contrast, low burden granulomas exhibit cell-cell interactions consistent with Type I immune responses (*CCL3, CXCL9/10/11, DLL1, IFNG, IL18*) and Th17 chemoattraction (*CXCL16, CCL20*) and successful immune mobilization and activation (Li et al., 2013; Lim et al., 2008; Touzot et al., 2014). Ligands specifically associated with low burden granulomas include co-stimulatory molecules important in immune activation (*CD40LG, CD48, CD70, CD80, CD86*), those involved in lymphocyte adhesion (*CD58*), and antimicrobial peptides (*DEFB1, SLPI*) (**Fig. 5I**) (Tateosian et al., 2012). Various antimicrobial peptides have been implicated previously in direct control of MTB infection (Fabri et al., 2011; Liu et al., 2006); whether intercellular communication is an essential or auxiliary role remains to be determined. Importantly, production of low burden-linked ligands is distributed across the cell types of Groups 1, 3, 4, and 5, but not Group 2 (**Fig. 5J**); signaling occurs between multiple T and macrophage cell subsets, suggesting that successful Mtb control requires coordinated interactions across diverse innate and adaptive immune cell types. Supporting this interpretation, gene set enrichment analyses on ligands and receptors whose interactions are strengthened in low burden granulomas revealed enrichment for processes including T cell activation and differentiation and signaling associated with pro-inflammatory cytokines (e.g., TNF) (**Fig S8A**). Likewise enriched in interactions associated with low burden granulomas are additional metabolic processes such as fatty acid metabolism and heat generation, which have been individually studied extensively in Mtb but here connect to broader signaling interactions associated with Mtb control.

Beyond ascribing a simple binary role to each cell type, our cell-cell interaction analyses also indicate context-dependent roles for particular cell types and ligands. For instance, with respect to cell types, the macrophage-dominated Group 1 is not statistically correlated with granuloma control in our compositional analyses (**Fig 5A**), but participates in the second most interactions in both high and low burden granulomas (**Fig 5B-C,E,G-H**). The idea of dual roles for Group 1 cells is borne out by examination of the ligands produced by Group 1 cell types in high (e.g., *PDGFB, CD5L, TNFSF13*) and low burden (e.g., *CXCL9/10/11, CD86, IL18, CCL20*) microenvironments (**Fig 5I-J**). Similarly, we observe that some individual ligands participate in interactions strengthened in both high and low burden granulomas, suggesting pleiotropic effects for these molecules. As one specific example, IL-1’s effects on Mtb control vary based on disease stage and model (Juffermans et al., 2000; Law et al., 1996; Mayer-Barber et al., 2014); based on our analyses, *IL1A* and *IL1B* each mediate interactions associated with both high and low bacterial burden, but are derived from different sender cell populations in the two instances. Thus, our intercellular interaction analyses uncover axes of cellular plasticity and ligand pleiotropy across granuloma microenvironments, important for improved understanding and therapeutic modulation of Mtb.

## Discussion

Within an individual with Mtb infection, distinct granulomas can achieve sterilizing immunity, immune standoff, or frank immune failure (Flynn, 2006; 2010; Lin *et al*., 2014b; Lin et al., 2009). In NHPs, which most closely recapitulate human Mtb infection and disease (Coleman et al., 2014a), this heterogeneity provides an opportunity to define the cellular and molecular factors that correlate with bacterial control to identify potential host-directed prevention and cure strategies for TB. While a spectrum of granuloma-level bacterial control has been appreciated previously, the immune correlates of bacterial control within granulomas have not been mapped comprehensively. By coupling advanced serial imaging, scRNA-seq, and molecular measures of bacterial growth and killing, the present study provides new insights into the immunologic control and temporal evolution of granulomas in Mtb infection: we discover and define how the timing of granuloma appearance correlates with distinct microenvironmental signaling networks formed through host responses and shapes eventual bacterial persistence or control. Overall, our data substantiate a model where the state of the surrounding host cellular ecosystem informs a granuloma’s infection trajectory, leading to long-term, stable states which either permit or restrict bacterial survival.

We find that high CFU, early granulomas are characterized by significantly higher proportions of mast cells and plasma cells, as well as a central group of cell types (further including fibroblasts, and endothelial cells) that exhibits extensive self-directed signaling exchanges. While mast cells have been described in granulomatous conditions, such as TB lymphadenitis (Taweevisit and Poumsuk, 2007), leprosy skin lesions (Bagwan et al., 2004), and liver granulomas (Celasun et al., 1992), and may orchestrate immune cross talk in TB (Garcia-Rodriguez et al., 2017), this is the first description of direct correlation with failure of Mtb control in TB granulomas. Structurally, we find mast cells inter-digited in the lymphocyte cuff of TB granulomas, physically well positioned to play significant regulatory roles.

The mast cells in high-burden granulomas are major producers of Type II cytokines, especially *IL4, IL5,* and *IL13*, which are important down-modulators of lymphocyte and macrophage antimicrobial activity, including inhibiting the cytolytic functions of CD8+ T cells (Kienzle et al., 2005; Wijesundara et al., 2013). However, IL4 and IL13 have broader functions in the context of wound healing. Indeed, the cellular interactions in high burden granulomas reveal both specific signaling molecules (e.g., FGF1 from Type 1 pneumocytes, PDGFB from endothelial cells, ANGPTL4 from plasma and mast cells, among others) and broad pathways (e.g., FGF and VEGF, among others) that reflect fibrosis, metabolic remodeling, and angiogenesis. Collectively, these data suggest a cascade of interactions in early appearing granulomas with failed control, whereby an initially permissive environment is reinforced by a tissue remodeling response that seeks to limit and wall off pathologic activity, thereby allowing for persistence of both Mtb and the Type II/wound healing microenvironment itself. While more detailed studies on the roles of wound healing responses and tissue remodeling in TB are indicated, these features may represent critical targets for host-directed therapies that not only need to enhance restrictive adaptive immune responses but also address the maladaptive features of microenvironments permissive to granuloma persistence.

Indeed, it is striking how strongly the timing of granuloma appearance (as identified by PET-CT imaging) correlates with the formation of distinct classes of complex yet stable cellular communities and their accompanying levels of bacterial control. We note that granulomas identified late by PET-CT imaging may either be formed later—for example through dissemination (Martin *et al*., 2017) —or take more time to reach the threshold to be identified by PET-CT scans (limit of detection ≥1mm) because of more efficient immune control or differences in the quality of the inflammatory response (Cronan et al., 2021). Regardless of the exact mechanism, late appearing granulomas are characterized by the enrichment of multiple T and NK cell subsets, as well as extensive pro-inflammatory, pro-activating, and pro-migratory signaling networks predominated by T cell subsets, which may exclude or prevent the establishment of self-reinforcing Type II signaling. Moreover, our measures of cumulative bacterial burden (CEQ) indicate that late granulomas have lower bacterial burden because of greater bacterial killing (CFU/CEQ), linking these adaptive immune features to true sterilizing immunity.

The strongest cellular correlate of bacterial control was a subcluster of cells with transcriptional features of both Type 1 and Type 17 T cells that was expanded in granulomas with bacterial control. Aspects of these data are consistent with recent observations that granulomas established in immune primed environments—e.g., existing Mtb infection (Cadena et al., 2018) or intravenous or intrabronchial BCG vaccination—are characterized by Th1/17 expression patterns that are associated with protection (Darrah *et al*., 2020; Dijkman et al., 2019); however, we extend these findings, defining appreciable substructure among the T1-T17 subcluster of relevance to control. The *CD4* T1-T17 subpopulation (subpopulation 1) is most consistent with published descriptions of Th1/17 cells (e.g., Th1* or ex-Th17) (Amezcua Vesely *et al*., 2019). These cells may represent precursors to long lived tissue memory, which has been shown to play a crucial protective role in autoimmunity, bacterial control, and memory immune responses to pathogens (Amezcua Vesely *et al*., 2019; Liang et al., 2015; van Hamburg and Tas, 2018; Wacleche et al., 2016), including Mtb infection. A recent study using flow cytometry and immunohistochemistry in Mtb infected rhesus macaques support an association of Th1 (IFNγ+) and Th17 (IL-17+) cells in lung tissue with latent infection (Shanmugasundaram et al., 2020); in contrast, another study using scRNA-seq reported activated CD4 and CD8 T cells including Th1 and Th17 in the lung tissue of macaques with pulmonary TB (Esaulova et al., 2021). The *CD8* subsets within the T1/T17 subcluster (subpopulations 2 & 3), meanwhile, have not been described previously. The former of these is strongly associated with bacterial control and may represent a novel immunologic paradigm that can be exploited for vaccine development. Subpopulation 3 intriguingly, expresses elevated *TNF* and *IFNG* but does not associate with bacterial restriction; further profiling will be necessary to establish the significance of this subset and its relation to previously appreciated Type 1 and Type 17 features of control (Algood *et al*., 2005; Gideon *et al*., 2015; Green *et al*., 2013; Khader *et al*., 2007; Khader and Gopal, 2010; Lin *et al*., 2007; Lyadova and Panteleev, 2015; Millington *et al*., 2007; O’Garra *et al*., 2013; Scriba *et al*., 2017).

Our data also revealed an interesting CD4 and CD8 expressing T cell subcluster associated with low burden granulomas that resembles stem-like T cells (Ahmed *et al*., 2016; Caccamo *et al*., 2018; Cartwright et al., 2016; Fuertes Marraco et al., 2015; Gattinoni *et al*., 2011; Mateus et al., 2015; Todryk, 2018). We hypothesize that these cells may be a source of T cell renewal in granulomas and may differentiate into the various functional subsets we observe within them. It is possible, however, that these represent memory T cells that are not specific for Mtb antigens but migrate to the granuloma in response to inflammation and/or chemokine gradients. Indeed, flow-cytometry based studies support that a majority of T cells in granulomas do not respond to Mtb antigens by making cytokines and do not display hallmarks of exhaustion (Gideon *et al*., 2015; Sakai et al., 2016; Wong *et al*., 2018). These stem-like T cells warrant additional study, as they associate with control of Mtb in granulomas and, if antigen specific, could be explored as a potential vaccine target.

Although both CD4 and CD8 T cells have been implicated in control of Mtb infection, the cytotoxic function of lymphocytes in Mtb infection has been relatively understudied, with emphasis placed instead on macrophage activating cytokines, such as IFN-γ and TNF. However, we also find previously unappreciated complexity among granuloma cytotoxic cells of relevance to bacterial control. In accordance with another recent study (Rath *et al*., 2020), our 6 cytotoxic T/NK subclusters do not align neatly with canonical markers of cellular identity that would define them as classical CD8αβ or CD4 T cells, NK, NK T cells, or γδ T cells, but instead appear to be variable mixtures of innate and adaptive cell types with common transcriptional programming. Of these, cytotoxic subcluster 4, which is enriched in CD8αβ T cells and defined by expression of several granzymes and perforin, likely represents cytotoxic effector T cells that target infected cells for apoptosis and is associated with low burden granulomas. A recent study on lung tissue from Mtb infected macaques also found evidence of cytotoxic molecule expression associated with controlled infection (Esaulova *et al*., 2021). These findings reveal the importance of cytotoxic innate and adaptive lymphocytes in temporal control of Mtb in granulomas, and a potential role for in future prevention and cure strategies.

Complementing analyses that characterize individual cell states associated with Mtb control, our cell-cell interaction analyses support connections between control, timing of granuloma appearance, and primed immune responses. Robust control of Mtb at the granuloma level correlated with interactions between particular subsets of T cells and macrophages and was mediated via specific proinflammatory cytokines (e.g., CCL3, IFNγ), T cell chemoattractants (e.g., CXCL9/10/11/16, CCL20), and co-stimulatory molecules (e.g., CD40LG, CD80, CD86). The pro-inflammatory T cell-mediated signaling in late-appearing, low bacterial burden granulomas stands in contrast to early type II immune and would healing activities in high bacterial burden granulomas, highlighting key cell types and interdependencies behind integrated, holistic host responses.

Importantly, our analyses reveal not just sets of biological pathways utilized in the host cells of high vs. low burden granulomas, but also assign roles to the specific cell types that drive these signaling patterns. In particular, the strong internal signaling among Group 2 cell types and comparatively weaker cross-talk to other groups in early lesions may drive establishment of a cellular ecosystem dominated by Type II immune and wound healing responses that preclude effective T cell engagement and conversion to a more restrictive state. By comparison, in late-appearing lesions, primed T cell populations, in concert with different innate populations, may use a variety of pro-inflammatory and pro-activation interactions to control Mtb growth or dissemination; a similar phenomenon might explain how infection with Mtb can protect against subsequent reinfection (Lin *et al*., 2014b), even in the presence of ongoing original infection, by locally recruiting adaptive responses that can act before self-reinforcing Group 2 responses work to limit pathology. Future work will be necessary to determine the relative importance of each adaptive response for control. More broadly, we will need to define the relative stability of these two broad cellular microenvironments and how host perturbations—whether vaccination, therapies or coinfections—impact their balance.

In addition to identifying cellular populations that relatively exclusively associate with high or low burden granulomas, we find cellular plasticity among some cell populations which appear capable of producing ligands linked to either bacterial persistence or control. For example, the Group 1 macrophage populations vary their interaction patterns, perhaps based on the signals they receive from their microenvironment. These responses may, in turn, help mold the phenotypes of surrounding cell types via an immunologic feedback loop (e.g., contributing to persistence of wound healing and Type II immune signaling, or to effective immune recruitment and activation for granuloma clearance). Indeed, we find that individual ligands (IL15, TNFSF13, IL1A and IL1B) can also exhibit pleiotropic effects and participate in interactions enriched in either high or low burden granulomas. Such pleiotropic ligand effects may arise from differing spatial contexts around sender cells (e.g., whether TNFSF13 is secreted from lymphocyte cuff-localized mast cells vs. from macrophage populations closer to the granuloma core), or from combinatorial interactions with other ligands whose presence varies with the distinct microenvironmental ecosystems of high vs. low burden granulomas. These results may help reconcile contrasting findings on ligands’ roles differing by disease stage and model (e.g., IL-1), but also inform the selection of targets for therapies seeking to unwind deleterious microenvironments or reinforce adaptive responses (Juffermans *et al*., 2000; Law *et al*., 1996; Mayer-Barber *et al*., 2014).

Importantly, we note that the contrasting microenvironments revealed through our analyses can occur within the same individual. This suggests the importance of rationally designing new classes of host directed TB preventions and cures that seek to destabilize one set of interactions while reinforcing the other. Indeed, the current standard of care for Mtb calls for multiple antibiotics to be administered for months and has largely remained unchanged for decades (Keshavjee and Farmer, 2012). Our work now defines the complexities of cellular ecosystems encapsulated as granulomas (e.g., reprogramming of plastic tissue-resident cells, recruitment of non-resident immune cells, etc.). Knowledge of intercellular networks underlying granuloma stability will spur future research efforts that identify and manipulate linchpins that serve as key nodes in limiting or enhancing the efficacy of therapeutic and prophylactic measures. For instance, the ligands and receptors implicated in interactions strengthened in high burden granulomas are also enriched for targets of several vitamin A derivatives, including alitretinoin, beta-carotene, and retinol (**Fig S8B**), consistent with vitamin A’s known activity in promoting wound healing. We note, however, that vitamin A deficiency is a strong risk factor for progression to TB disease (Albana et al., 2017). These observations may be consistent with wound healing responses that create conditions in which bacteria cannot be eradicated but can be contained, speaking to the complexity of intervening in Mtb pathology. In contrast, low burden-linked signaling molecules are also enriched for targets of immunomodulatory drugs used to treat dermatoses and keratoses (e.g., fludroxycortide, imiquimod) (**Fig S8B**), aligning with a model where successful immune activation circumvents the need for wound healing responses. To most effectively target these complex granuloma ecosystems, we will need new computational methods that can pinpoint the relative importance of different molecular targets and cell types to granuloma stability and determine the most promising points of intervention to destabilize and modulate a densely interacting multicellular community toward adaptive states.

To fully optimize these host directed therapies, additional work on intercellular communication will be necessary since certain classes of regulatory and effector interactions are not fully captured in this type of analysis. For example, as part of our cell-cell interaction analyses, we found strong enrichment for the expression of distinct neuro-hormonal modulators by Group 2 (e.g., *NRG1, RLN3, NTS*) and Group 3 cells (e.g., *NRG2, UCN3*) but did not capture any potential neural interactors in our scRNA-seq dataset, limiting our ability to discern fully how they associate with, and might be leveraged to achieve, control. Nonetheless, ligands and receptors implicated in low-burden interactions are enriched for targets of several neuropsychiatric agents, including spiperone, scopolamine and serotonin, where serotonin reuptake inhibitors have already been identified in screens for host-acting compounds that improve macrophage control of Mtb, supporting potential for their further investigation (Heemskerk et al., 2021; Stanley et al., 2014) . Equally critically, a significant proportion of cell types in Group 3 expressed cytolytic effector genes that can directly drive bacterial control, suggesting a potential therapeutic role for IL15 super-agonists in clinical development that can drive expansion of cytotoxic populations.

It should be noted that granulomas are inherently heterogenous and include necrotic debris, requiring robust technical correction and quality control; this results in an analysis of only high-quality cells. Since only a fraction of cells from each granuloma are analyzed, proportions may not reflect the true composition of cells within a granuloma and may be skewed toward lymphocytes, highlighting the importance of orthogonal validations. In bulk RNA-sequencing analysis of a distinct set of early and late granulomas, we observe generally similar trends in cell-type composition, supporting our conclusions. Relatedly, the transcriptomic granuloma landscape investigated here is from a single (albeit pivotal) time point, while including granulomas across a spectrum of growth trajectories. It is likely that expression of certain genes that arise early in infection and then are downregulated as infection progresses will be missed, as will some populations critical to guiding overall granuloma outcome. More generally, matched analyses of earlier and later time point post-infection, along with analysis of lung tissue and granulomas from vaccinated or reinfected and protected animals, will provide a more complete picture of the temporal control of Mtb in granulomas and is the subject of future work.

In summary, our data represent the first scRNA-seq investigation of the cellular and molecular features that dynamically associate with natural control of Mtb in pulmonary granulomas. Beyond recapitulating canonical correlates, our analysis defines nuanced actionable innate and adaptive functional cell states, and sheds light on essential dynamics among host-pathogen interactions (Iwasaki and Medzhitov, 2015). Collectively, our data substantiate a model where high Mtb burden within granulomas is dictated at a local level by Type II immune and tissue-protective (wound healing) responses that seek to maintain essential tissue functionality, at the expense of creating a niche for bacterial persistence. In granulomas that form later in infection and therefore in the context of an adaptive immune response, this balance is tipped towards bacterial control by the emergence of adaptive T1-T17 and cytotoxic responses, with interactions involving innate immune cell types enabling sufficient infiltration and activation of these T cell subsets. As a result, successful immune coordination across cell types in late-forming granulomas may obviate the self-reinforcing Type II/wound healing response that would otherwise exclude immune effector functions needed for Mtb control. We also identify cell types and ligands that participate in both high and low burden granulomas, potentially indictive of phenotypic plasticity and pleiotropic effects that may both be molded by and (in turn) reinforce distinct, pathology-associated granuloma microenvironments. Such a framework is consistent with previous observations of natural (Cadena *et al*., 2018) or induced (Darrah *et al*., 2020) control, and supports the need to look to new combinatorial host-directed paradigms for the development of novel efficacious therapeutic and prophylactic measures. Moving beyond the perspective of individual molecular targets, our work highlights the importance of the complexities of divergent host cellular ecosystems in driving Mtb persistence or control. By defining and nominating several putative axes of intra-and intercellular signaling associated with contrasting Mtb outcomes, our work provides a foundation for enabling effective manipulation of the properties and states of complex cellular ecosystems, therapeutically-relevant destabilization of pathologic molecular environments to enable adaptive immune access, and fundamental connections to other inflammatory and infectious diseases that affect epithelial barrier tissues (Hughes et al., 2020; Ordovas-Montanes *et al*., 2018).

## Acknowledgments

We are grateful to the research and veterinary technicians: Chelsea Chedrick, Carolyn Bigbee, Nicholas Schindler, Mark Rogers, Tara Rutledge, Chelsea Causgrove and Brianne Stein in the Flynn lab who assisted with this work, as well as helpful discussions with members of the Flynn, Scanga, Mattila, Lin and Shalek laboratories. We also thank the efforts of the University of Pittsburgh Division of Laboratory Animal Research technicians for husbandry of the animals.

## Funding

Bill and Melinda Gates Foundation (OP1139972: AL, SMB, SMF, JLF, AKS; OPP1202327: AKS)

Searle Scholars Program (AKS)

The Beckman Young Investigator Program (AKS)

Sloan Fellowship in Chemistry (AKS)

NIH (5U24AI118672, BAA-NIAID-NIHAI201700104) (AKS)

American Lung Association RG571577(HPG)

F30-AI143160 (TKH) NIH T32A1065380 (NLG)

NSF GRFP grant (CNT, SKN 1122374)

Fannie and John Hertz Foundation Fellowship (CNT)

Wellcome Trust Fellowship award 210662/Z/16/Z (AL)

Koch Institute Support (core) grant P30-CA14051 from the National Cancer Institute (CL)

NIH CFAR P30 AI060354 (BB)

NIH R01A1022553 (BB)

T32 A1007387 (JR)

NIH K12 (CW)

## Author contributions

Conceptualization: JLF, SMF, AKS

Data Curation: HPG, TKH, FFH, PM, AGW, NLG, AL

Formal Analysis: HPG, TKH, CNT, NLF, FFH, AGW,

Methodology: HPG, TKH, MHW, CNT, AAT, TMG, FFH, CK, PM, AGW, SKN, HJB, BB, JCL

Investigation: HPG, TKH, MHW, CNT, AAT, TG, FFH, JW, CK, JMP, PM, AGW, SKN, HJB, SKCG, AM, KVP, CLA, CTC, JAT, LJF, HJB, PLL, SI, JYP, JMR, AS, JOM

Visualization: HPG, TKH, CNT

Validation: HPG, TKH, NLG, KN, CGW, SI

Resources: JLF, SMF, AKS, JCL. RM, AL

Funding acquisition: JLF, SMF, AKS

Project administration: CAS

Supervision: JLF, SMF, AKS, SMB, BDB, AL, JCL, BB

Writing – original draft: HPG, TKH, SMB, JLF, SMF, AKS

Writing – review & editing: HPG, TKH, CNT, MC, SMB, JLF, SMF, AKS

## Data and materials availability

### Lead Contact

Further information and requests for resources and reagents should be directed to and will be fulfilled by the Lead Contact, Alex K. Shalek.

### Materials Availability

The study did not generate new unique reagents.

### Data and Code Availability

Raw and processed data will be available on the Gene Expression Omnibus and can be accessed at https://singlecell.broadinstitute.org/single_cell/study/SCP257/cellular-ecology-of-m-tuberculosis-granulomas?scpbr=the-alexandria-project#study-visualize. Additional code is available upon request from the lead contact.

## Materials and Methods

### Ethics Statement

All experimental manipulations, protocols, and care of the animals were approved by the University of Pittsburgh School of Medicine Institutional Animal Care and Use Committee (IACUC). The protocol assurance number for our IACUC is D16-00118. Our specific protocol approval numbers for this project are 18124275 and IM-18124275-1. The IACUC adheres to national guidelines established in the Animal Welfare Act (7 U.S.C. Sections 2131 - 2159) and the Guide for the Care and Use of Laboratory Animals (8^th^ Edition) as mandated by the U.S. Public Health Service Policy.

All macaques used in this study were housed at the University of Pittsburgh in rooms with autonomously controlled temperature, humidity, and lighting. Animals were singly housed in caging at least 2 square meters apart that allowed visual and tactile contact with neighboring conspecifics. The macaques were fed twice daily with biscuits formulated for nonhuman primates, supplemented at least 4 days/week with large pieces of fresh fruits or vegetables. Animals had access to water *ad libitem*. Because our macaques were singly housed due to the infectious nature of these studies, an enhanced enrichment plan was designed and overseen by our nonhuman primate enrichment specialist. This plan has three components. First, species-specific behaviors are encouraged. All animals have access to toys and other manipulata, some of which will be filled with food treats (e.g. frozen fruit, peanut butter, etc.). These are rotated on a regular basis. Puzzle feeders foraging boards, and cardboard tubes containing small food items also are placed in the cage to stimulate foraging behaviors. Adjustable mirrors accessible to the animals stimulate interaction between animals. Second, routine interaction between humans and macaques are encouraged. These interactions occur daily and consist mainly of small food objects offered as enrichment and adhere to established safety protocols. Animal caretakers are encouraged to interact with the animals (by talking or with facial expressions) while performing tasks in the housing area. Routine procedures (e.g. feeding, cage cleaning, etc) are done on a strict schedule to allow the animals to acclimate to a routine daily schedule. Third, all macaques are provided with a variety of visual and auditory stimulation. Housing areas contain either radios or TV/video equipment that play cartoons or other formats designed for children for at least 3 hours each day. The videos and radios are rotated between animal rooms so that the same enrichment is not played repetitively for the same group of animals.

All animals are checked at least twice daily to assess appetite, attitude, activity level, hydration status, etc. Following *M. tuberculosis* infection, the animals are monitored closely for evidence of disease (e.g., anorexia, weight loss, tachypnea, dyspnea, coughing). Physical exams, including weights, are performed on a regular basis. Animals are sedated prior to all veterinary procedures (e.g. blood draws, etc.) using ketamine or other approved drugs. Regular PET/CT imaging is conducted on most of our macaques following infection and has proved very useful for monitoring disease progression. Our veterinary technicians monitor animals especially closely for any signs of pain or distress. If any are noted, appropriate supportive care (e.g. dietary supplementation, rehydration) and clinical treatments (analgesics) are given. Any animal considered to have advanced disease or intractable pain or distress from any cause is sedated with ketamine and then humanely euthanatized using sodium pentobarbital.

### Research Animals

Four Cynomolgus macaques (*Macaca fascicularis*), >4 years of age, (Valley Biosystems, Sacramento, CA) were housed within a Biosafety Level 3 (BSL-3) primate facility as previously described and as above. Animals were infected with low dose (∼10 colony-forming units (CFUs)) *M tuberculosis* (Erdman strain) via bronchoscopic instillation. Infection was confirmed by PET-CT scan at 4 weeks and monitored with clinical and radiographic examinations until 10 weeks post infection.

### Serial PET-CT Imaging

Animals underwent PET-CT scans after Mtb infection at 4 weeks, 8 weeks and pre necropsy (i.e. 10 weeks post-infection) as previously described (White *et al*., 2017). Briefly, animals were sedated, intubated and imaged by 2-deoxy-2-^18^F-D-deoxyglucose (FDG) PET imaging (microPET Focus 220 preclinical PET scanner, Seimens Medical Solutions, USA, Malvern, PA) and Cretom CT scanner (Neurologica Corp, Danvers, MA, USA) within biosafety level 3 facility. The total lung FDG avidity was analyzed using Osirix viewer, an open-source PACS workstation and DICOM viewer (Pixmeo, Bernex, Switzerland). The whole lung was segmented on CT by using the growing region algorithm on the Osirix viewer to create a ROI of normal lung (Hounsfield units < 200). The closing tool was used to include individual nodules and other pulmonary disease. The ROI was transferred to the co-registered PET scan and manually edited to ensure all pulmonary disease was included. Voxels outside the ROI were set to zero and voxels with an SUV greater than or equal to normal lung (SUV > 2.3) were isolated. Finally, the “Export ROIs” plug-in was then used to export the data from these isolated ROIs to a spreadsheet where the total SUV per voxel were summed to represent the total lung FDG activity. Total FDG activity in lungs was used to estimate thoracic bacterial burden prior to reinfection (**Figure 1C**), as previously published(Coleman *et al*., 2014b; White *et al*., 2017). Granulomas were individually characterized by their date of establishment (scan date), size (mm), and relative metabolic activity as a proxy for inflammation ([^18^F]-FDG standard uptake normalized to muscle [SUVR])(Coleman *et al*., 2014b; White *et al*., 2017). Granulomas greater than 1mm are detected by CT scan.

### Necropsy

Necropsy was performed as previously described (Gideon *et al*., 2015; Lin *et al*., 2013; Lin *et al*., 2009; Maiello *et al*., 2018). Briefly, an ^18^F-FDG PET-CT scan was performed on every animal 1-3 days prior to necropsy to measure disease progression and identify individual granulomas. At necropsy, monkeys were maximally bled and humanely sacrificed using pentobarbital and phenytoin (Beuthanasia; Schering-Plough, Kenilworth, NJ). Individual granulomas previously identified by PET-CT and those that were not seen on imaging from lung and mediastinal lymph nodes were excised for histological analysis, bacterial burden, and other immunological studies. TB specific gross pathologic lesions and overall gross pathologic disease burden was quantified using a previously published method (Maiello *et al*., 2018). The size of each granuloma was measured by pre-necropsy scans and at necropsy. Granulomas were enzymatically dissociated using the Gentlemacs dissociator system (Miltenyi Biotec Inc) to obtain single cell suspension and used to enumerate bacterial burden and applied on a Seq-Well device for single cell RNA-sequencing (scRNA-seq).

### Bacterial burden

200 μl of each granuloma homogenate were plated in serial dilutions onto 7H11 medium, and the CFU of *M. tuberculosis* growth were enumerated 21 days later to determine the number of bacilli in each granuloma (Gideon *et al*., 2015). As a quantitative measure of overall bacterial burden, a CFU score was derived from the summation of the log-transformed CFU/gram of each sample at the time of necropsy.

### Chromosomal equivalents, CEQ

DNA extraction and qPCR was performed with modifications as described previously ((Lin *et al*., 2014b)). Briefly, frozen aliquots of homogenates were thawed and volumes recorded throughout the extraction process. Samples were transferred to tubes containing 150 μl of 0.1mm zirconia-silica beads (Biospec Products) before adding 600μl of Tris-EDTA buffer, pH 8.0. Three hundred microliters of phenol/chloroform/isoamyl alcohol (25:24:1, Sigma-Aldrich) at 70°C were subsequently added and the samples incubated at room temperature for 10 minutes. The samples were then vortexed, the aqueous layer separated and supplemented with 50 μl 5M NaCl and a second phenol chloroform extraction performed on the extracted aqueous layer. DNA was precipitated with the addition of one volume of 100% isopropanol and one-tenth volume of 3M sodium acetate and incubating at -20°C overnight. The DNA pellet was washed with 70% ethanol, dried and resuspended in nuclease-free water. Mtb genomes were then quantified using Taqman Universal Master Mix II (Life Technologies) and previously published *sigF* primer-probe combination (Lin *et al*., 2014b). Each sample was amplified in triplicate using an ABI Systems 7900HT machine. Chromosomal equivalents (CEQ) were quantified by comparing the samples with a standard curve derived from serial dilution of Mtb genomes prepared from liquid culture. Our detection limit for the standard curve was 10 copies per reaction. When we calculated the number of genomes for the whole granuloma, our detection limit was 1,000 copies per granuloma. Of the 26 granulomas analysed, 2 granulomas failed at the CEQ quantification and they were eliminated from CEQ and CFU/CEQ analysis.

### Immunohistochemistry analysis

Granulomas from macaques were harvested at 10 or 11 weeks post Mtb infection from other published (Phuah *et al*., 2016) and unpublished studies at the University of Pittsburgh. Following formalin fixation and paraffin embedding, 5 µm sections were placed on slides for staining. Slides were deparaffinized in xylenes, hydrated in a series of graded ethanol dips, and then antigen retrieval was performed by boiling the slides in a pressure cooker containing antigen retrieval citrate buffer for slides stained with c-kit and tryptase or Tris-EDTA buffer (Mattila *et al*., 2013) for slides stained with CD11c, CD20, and CD3. Sections were cooled to room temperature and washed with 1X PBS then stained overnight at 4°C in a humidified chamber using anti-human c-kit , anti-mast cell tryptase antibodies, or rabbit-anti-CD3 and mouse anti-CD11c antibodies as previously described (Phuah *et al*., 2016). For the c-kit and tryptase stained slides, the tissue sections were washed three times using 1X PBS and then incubated with anti-mouse IgG1 AF546 to label the anti-c-kit antibodies for 1 hour at room temperature in a humidified chamber. Tryptase staining was performed overnight at 4°C with anti-tryptase antibodies that were labeled with an Alexa Fluor 488 anti-rabbit IgG Zenon labeling kit. For the CD3, C11c, and CD20 stained sections, the CD3 and CD11c antibodies were labeled with donkey anti-rabbit IgG Alexa Fluor 647 and anti-mouse IgG Alexa Fluor 488-conjugated secondaries purchased Jackson ImmunoResearch Laboratories (West Grove, PA) or ThermoFisher, respectively. After the secondary antibodies were removed with PBS washes, CD20 was stained with rabbit anti-CD20 that was labeled with Alex Fluor 546 anti-rabbit IgG Zenon labeling kit. For both staining panels, the sections were washed again in 1X PBS and coverslips were applied using ProLong Gold Antifade Mountant with DAPI. For the slides stained with CD3, CD11c, and CD20, individual image channels were acquired with an Olympus FluoView 500 laser scanning confocal microscope (Olympus, Life Sciences Waltham, MA) maintained by the University of Pittsburgh’s Center for Biologic Imaging and combined and pseudocolored with the FIJI build of ImageJ (Schindelin et al., 2012). Images of c-kit and tryptase-stained slides were acquired with a Nikon e1000 epifluorescence microscope (Nikon Instruments, Melville, NY) operated by the NIS-Elements AR software package (Nikon).

Human granulomas were identified from sections of lung tissue obtained at subjects undergoing partial lung resection for clinical indications at King Dinzulu Hospital and Inksosi Albert Luthili central Hospital in Durban, South Africa. Gross pathology was assessed by Haematoxylin and Eosin (H&E) staining. Briefly, samples of lung were fixed in 10% neutral buffered formalin and processed routinely in a vacuum filtration processor using a xylene-free method with isopropanol as the main substitute fixative. Tissue sections were embedded in paraffin wax. Sections cut at 4 µm using a microtome, heated at 56°C for 15 min, dewaxed through two changes of xylene and rehydrated through descending grades of alcohol to water and stained with Haematoxylin & Eosin (H&E, 5 minute incubation with each stain). Slides were dehydrated in ascending grades of alcohol, cleared in xylene, and mounted with a mixture of distyrene, plasticizer, and xylene (DPX). For immunohistochemistry, 4 µm sections and were mounted on charged slides and heated at 56°C for 15 min. Mounted sections were dewaxed in xylene followed by rinsing in 100% ethanol and 1 change of SVR (95%). Slides were then washed under running water for 2 min followed by antigen retrieval via Heat Induced Epitope Retrieval (HIER) in Tris-sodium chloride (pH 6.0) for 30 minutes. Slides were then cooled for 15 min and rinsed under running water for 2 min. Endogenous peroxide activity was blocked using 3% hydrogen peroxide for 10 min at room temperature (RT). Slides were then washed in phosphate-buffered saline with 1% Tween (PBST) and blocked with protein block (Novolink) for 5 min at RT. Sections were incubated with primary antibodies for CD117 (A4502-CD117,c-kit, DAKO, 1:500), followed by washing and incubation with post primary (Novolink) for 30 minutes at RT. Slides were washed with PBST followed by incubation with the polymer (Novolink) for 30 min at RT. Slides were then washed and stained with DAB for 5 min, washed under running water and counterstained with hematoxylin for 2 min. Slides were rinsed under running water, blued in 3% ammoniated water for 30 s, washed under water, dehydrated and mounted in DPX.

### Flow cytometry

Granulomas harvested from other Mtb infected NHPs were used in the flow cytometery analysis and processed as previously published(Gideon *et al*., 2015). Cells were counted and stained for viability using fixable viability dye (Zombie NIR, Biolegend) and other surface and intracellular markers using the standard protocols. Surface markers include: CD3 (SP34-2, BD), CD4 (L200, BD), CD8a (RPA-T8, BD), CD8b (2ST8.5H7, BD), TCR gd (5A6.E9, ThermoFisher), CD16 (3G8, BD), NKG2A (Z199, Beckman Coulter and intracellular markers include: Granzyme B (GB11, BD), Granzyme A (CB9, BD) and Granzyme K (G3H69, BD). Samples were acquired on a Cytek Aurora spectral cytometer (5 laser configuration) and unmixed using SpectroFlo software (Cytek). Final analysis was performed in FlowJo (v10, FlowJo)

### Single-cell RNA-Sequencing (scRNA-seq)

High-throughput scRNA-seq was performed using the Seq-Well platform as previously described (Gierahn *et al*., 2017). Briefly, total cell counts from single-cell suspension of granuloma homogenate were enumerated and ∼15,000-30,000 cells were applied to the surface of a Seq-Well device loaded with capture beads in the BSL-3 facility at University of Pittsburgh. Following cell loading, Seq-Well devices were reversibly sealed with a polycarbonate membrane and incubated at 37°C for 30 minutes. After membrane sealing, Seq-Well devices were submerged in lysis buffer (5 M guanidine thiocyanate, 10 mM EDTA, 0.1% β-mercaptoethanol, 0.1% Sarkosyl) and rocked for 30 minutes. Following cell lysis, arrays were rocked for 40 minutes in 2 M NaCl to promote hybridization of mRNA to bead-bound capture oligos. Beads were removed from arrays by centrifugation and reverse transcription was performed at 52°C for 2 hours. Following reverse transcription, arrays were washed with TE-SDS (TE Buffer + 0.1% SDS) and twice with TE-Tween (TE Buffer + 0.01% Tween20). Following ExoI digestion, PCR amplification was performed to generate whole-transcriptome amplification (WTA) libraries. Specifically, a total of 2,000 beads were amplified in each PCR reaction using 16 cycles as previously described (Gierahn *et al*., 2017). Following PCR amplification, SPRI purification was performed at 0.6x and 0.8x volumetric ratios and eluted samples were quantified using a Qubit. Sequencing libraries were prepared by tagmentation of 800 pg of cDNA input using Illumina Nextera XT reagents. Tagmented libraries were purified using 0.6x and 0.8x volumetric SPRI ratios and final library concentrations were determined using a Qubit. Library size distributions were established using an Agilent TapeStation with D1000 High Sensitivity ScreenTapes (Agilent, Inc., USA).

### Bulk RNA Sequencing

Bulk RNA sequencing was performed using cells obtained from a total of 12 granulomas from a separate set of animals infected with Mtb for 10 weeks. Initially, granulomas were enzymatically dissociated and cells from each granuloma were placed in 100 uL of lysis buffer. RNA was then extracted from whole lysates using RNEasy kits (Qiagen, Inc.) and combined with mRNA capture beads. Reverse transcription, whole transcriptome amplification, tagmentation and sequencing were performed as described above. Within each bulk RNA sequencing sample, expression values were summarized across bead barcodes to obtain an aggregate expression profile for each population.

### Sequencing and Alignment

Libraries for each sample were sequenced on a NextSeq550 75 Cycle High Output sequencing kit (Illumina Inc., Sunnyvale, CA, USA). For each library, 20 bases were sequenced in read 1, which contains information for cell barcode (12 bp) and unique molecular identifier (UMI, 8bp), while 50 bases were obtained for each read 2 sequence. Cell barcode and UMI tagging of transcript reads was performed using DropSeqTools v1.12 (Macosko et al., 2015). Barcode and UMI-tagged sequencing reads were aligned to the Macaca fascicularis v5 genome (https://useast.ensembl.org/Macaca_fascicularis/Info/Index) using the STAR aligner. Aligned reads were then collapsed by barcode and UMI sequences to generate digital gene expression matrices with 10,000 barcodes for each array.

## QUANTIFICATION AND STATISTICAL ANALYSIS

### Data Processing and Quality Control

Initially, after examining a range of cell inclusion thresholds, a combined dataset of 169,830 barcodes was generated by applying a cutoff of 500 genes and 750 transcripts (UMIs). We visualized cells from each array using t-SNE across 30 principal components and performed Louvain clustering in Seurat. For many arrays, large clusters of cell barcodes were identified that were not marked by distinct cell-type defining gene expression. Instead, these cells were marked by distributed, low-level expression of genes presumed to originate from other cell types (e.g. *HBB* from erythrocytes, *JCHAIN* from plasma cells, and *CPA3* from mast cells). To understand the identity of these barcodes more fully, sequencing quality metrics were initially examined, and non-descript clusters did not significantly differ in the total number of aligned reads, detected genes, UMIs/cell, or mitochondrial percentage.

To more fully understand the identity of these clusters, multiple modeling approaches were pursued (**Figure S2**):

1. Initially, low-quality clusters were modelled as array-specific doublets. Here, models were constructed in which pseudo-doublets/multiplets (n=2, 5, 10, 15, or 20 cells) were created from random sampling of the remaining cell type clusters. However, in these models, there was not significant overlap between the generated pseudo-multiplets and the clusters with non-distinct gene expression patterns.
2. Random cells were created by binomial sampling a pseudo-population average expression vector generated by summation of expression profiles across all cell type clusters not suspected to be derived from ambient contamination. In these models, direct overlap was not observed between the simulated mixed population and those clusters with non-distinct gene expression patterns.
3. Finally, we examined whether these clusters might represent deep sampling of ambient contamination or cellular debris by generating a “contamination” scoring scheme. First, to identify the clusters within each array, 30 principal components were calculated (this was observed to consistently capture the majority of variation in each array), and Louvain clustering (resolution = 1.25) was performed using all significant principal components (JackStraw Empirical P-value < 0.05). Next, within each array, cluster-specific “contamination” scores were generated that consisted of 3 components:

a. **A measure of array-specific background contamination by cluster (“soup expression”).** For each array, a background expression profile was generated based on low-UMI barcodes (See *Correction for Residual Background Contamination* below for full details). A set of “soup”-defining genes was identified at a range of thresholds for soup-defining gene expression (0.01, 0.005, 0.001, and 0.0005;), a value that represents the proportional contribution of a given gene to the cumulative soup expression profile for each array. Array-specific, background-contamination scores were generated for the set of soup-defining transcripts using the AddModuleScore function in Seurat. Clusters with ambiguous/overlapping expression of lineage-defining gene expression signatures (Erythrocytes: *HBB*, Plasma cells: *JCHAIN*, Mast cells: *CPA3*, etc.) were observed to be significantly enriched for soup-defining gene expression. Finally, to calculate “contamination’ scores, expression scores for soup genes at a threshold of 0.001 were generated to calculate the average soup-profile score for each cluster within each array.
b. **An estimate of biological signal (“biological signal”)**. Here, the average log-fold change for the top 5 genes enriched within each cluster was calculated. For clusters dominated by ambient RNA, lower fold change enrichments for their biological signature genes were observed relative to clusters characterized by expression of canonical cluster-defining genes. In cases where the highest average log-fold change values within a cluster were below the “return threshold” in Seurat, we set the value to the default return threshold of 0.25.
c. **A measure of co-expression of lineage-defining genes (“soup linage coexpression”)**. 5 genes were manually selected that were recurrently over-represented in clusters suspected to arise from ambient contamination and cellular debris. Specifically, the following genes were selected: *HBB* (An erythrocyte-defining gene), *JCHAIN* (A plasma cell defining gene), *COL3A1* (A fibroblast defining gene), *SFTPC* (A type 2 pneumocyte defining genes), and *CPA3* (A mast cell defining gene). For each cell barcode, the number of these five genes with non-zero expression was calculated as a measure of lineage-defining co-expression. Within each cluster, the average co-expression of these genes was calculated and one was subtracted from this average to allow for endogenous expression of 1 lineage-defining gene. This parameter was specifically added to avoid exclusion of *bona fide* cell clusters with high-background contamination (presumably due to low endogenous RNA content) and low biological signal (e.g., naïve T cells). Here, cell populations that scored high for markers of a single lineage yet had higher soup-expression scores presented with lower rates of co-expression of these soup and lineage defining transcripts relative to clusters which did not, likely representing ambient RNA and debris.

Using these three values, cluster-specific background “contamination” scores were calculated for each array in 2 ways:

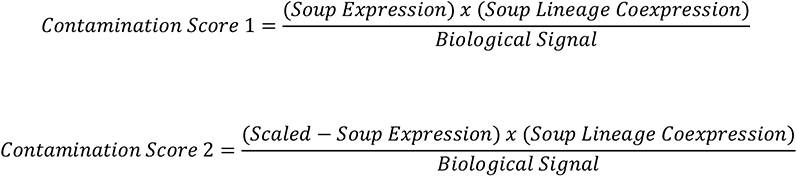

These two “contamination” scores quantify both the (1) absolute and (2) relative soup-profile contamination in subsequent cluster classification.

Next, for each array, clustering was performed to identify clusters with array-specific ambient contamination and debris. Specifically, hierarchical clustering was performed using a total of 7 variables to identify clusters defined by ambient contamination: the 2 contamination scores (shown above), three scaled soup scores (soup gene thresholds: 0.01, 0.05 and 0.001), the average log-fold change for the top 5 cluster genes, and soup/lineage gene co-expression. For each array, the hierarchical clustering tree was cut at the first branch point to identify clusters with a signature of ambient contamination. In total, 41 array-specific clusters, comprising 56,590 barcodes from 21 out of 32 total arrays, were identified as characterized by ambient RNA contamination and cellular debris and removed them in all subsequent analyses.

### Correction for Residual Background Contamination

After removal of cell barcodes that were derived from background contamination and extracellular debris, additional correction for ambient RNA contamination was performed among remaining cell barcodes on an array-by-array basis. Among filtered cell barcodes, array-specific, ambient RNA contamination was observed to be marked by ectopic expression of cell-type defining genes (e.g. widespread expression of *JCHAIN*, *HBB*, and *CPA3* etc.). Specifically, this contamination was observed to vary in relation to the overall distribution of cell types recovered from each array. To correct for residual ambient contamination within each array, SoupX (Young, 2018) was used to: (1) generate array-specific profiles of background contamination, (2) estimate per-cell contamination fractions, and (3) generate corrected background-corrected UMI counts matrices. To generate background expression profiles, counts matrices containing up to 50,000 barcodes were generated to assemble a collection of low-UMI cell barcodes that presumably represent extracellular mRNA. For each array, a UMI threshold for background expression was determined using EmptyDrops (Lun et al., 2019) to estimate the likelihood distribution that low-UMI barcodes represent cells rather than ambient contamination. Using an array-specific UMI-threshold (Range: 20-100 UMIs), a composite background profile was created for each array. To estimate the per-cell contamination fraction, a set of lineage-defining genes was first identified with bimodal expression patterns across cells (i.e., lineage defining genes with leaky expression). For each array, this set of soup-defining, lineage genes was used to estimate contamination fraction for cell types with known endogenous expression. Finally, the composite soup profile was subtracted from each the transcriptional profile of each cell based on the estimated contamination fraction. For each array, individual transcripts most likely to be contamination were removed from each single-cell based on the estimated contamination fraction. Specifically, individual transcripts were sequentially removed from each single-cell transcriptome until the probability of subsequent transcripts being soup-derived was less than 0.5 to generate a background-corrected counts matrix for each array **(Table S2b)**.

### Separation of Doublets

Within each array, doublet identification and separation were performed using DoubletFinder. To account for differences in cell loading densities and expected cell doublet frequencies, array-specific estimates of the expected number of doublets were generated **(Table S2a).** For example, for a total of 20,000 cells applied to a Seq-Well device containing 85,000 wells (lambda = 20,000), an expected doublet rate of >2.37% (since not all of the array’s surface area contains wells) was calculated. For each array, pseudo-doublets were generated using DoubletFinder (McGinnis et al., 2019). Here, the pK parameter estimate was separately optimized for each array by performing a parameter sweep in which we selected the pK value with the maximum bimodality coefficient, while a pN = 0.25 was maintained across all arrays based on published recommendations (McGinnis *et al*., 2019). Cells were identified as doublets based on their rank order in the distribution of the proportion of artificial nearest neighbors (pANN). Specifically, the pANN value for the cell at the expected doublet percentile was identified and the corresponding pANN value was used as a threshold to remove additional cells in the event of ties. In total, we excluded 3,656 cells as doublets (**Table S2a,c**).

### Integrated Cell Type Classification

Following the aforementioned quality filtering, a combined dataset of 109,584 cells was used in downstream analysis **(Table S2d)**. An initial dimensionality reduction was performed on these cells by selecting 1580 variable genes, performing principal component analysis (PCA), UMAP dimensionality reduction and Louvain clustering using Scanpy (Wolf et al., 2018). To identify broad cell types, we examined cluster assignments at multiple levels of clustering resolution (Resolutions: 0.5 to 2.25). We selected a cluster resolution of 1.00 because this was the resolution beyond which branching did not result in discovery of clusters that represent distinct cell lineages (e.g., division of Type 1 and Type 2 pneumocytes) (**Table S5**). To define major cell populations, extensive comparisons to existing signatures of lung parenchyma and immune cell populations were performed using data from the Tabula Muris (Tabula Muris *et al*., 2018) and Mouse Cell Atlas (Han *et al*., 2018) studies. Specifically, lung scRNA-seq data from both studies were collected and used to calculate enriched gene expression signatures for each lung cell type cluster using a Wilcox rank-sum test. For each cluster, the top 20 genes **(Table S3)** were selected as a cluster-specific expression signature and then used them to score all cells in the granuloma dataset. The average signature score within each cluster was calculated and the distribution of signature score was examined within each granuloma cell type, and significance was determined via permutation testing.

### Cell Type Assignment of Proliferating Cells

Among our top-level clusters was one defined by markers of cellular proliferation (*MKI67*, *TOP2A*, and *CDK1*). To identify the underlying cell type identity for these cells, a separate dimensionality reduction and clustering was performed among 3,123 cells defined by this proliferation signature. UMAP dimensionality reduction and Louvain clustering was running at multiple clustering resolutions (0.4-0.8), and a resolution of 0.70 was selected as the value beyond which no additional major cell type clusters were observed (**Figure S3E**). For each of the major cell types identified in the global clustering analysis, we generated a gene signature using the top 20 enriched genes and scored the proliferating cells clusters using the AddModuleScore function in Seurat. We then examined the distribution of cell-type signature scores across each of the sub-clusters of proliferating cells and re-assigned clusters based on enrichment of lineage-specific gene expression. Here, we assessed the significance of the cluster scores using a permutation test. More specifically, 1,000 permutations were performed in which the proliferating clusters were down-sampled to have the same number of cells. Cluster assignments of the cells were randomized and the average generic cell type signature score was calculated for each randomized cluster. The significance of a cell type score for each proliferating cluster was determined by comparing the observed average signature score to the random null distribution. Through this approach, distinct clusters of proliferating B cells, macrophages, neutrophils, plasma cells, and T cells were identified and re-assigned to their respective cell types.

### Filtering of Soup-Defining Transcripts

To avoid artifacts from ambient RNA contamination and cellular debris in sub-clustering of T cells and macrophages, genes that were observed to be soup-defining for any array were excluded. Specifically, a set of 210 soup defining genes was identified that comprised 0.001 of total soup expression in any array. The threshold of 0.001 was selected to maximize the cumulative fraction of soup expression with the least number of genes to avoid removing underlying biology. Here, this threshold value represents cumulative fraction of soup expression accounted for by a given gene for each array. In a further effort to avoid removing cell type specific biology, any genes with average log-fold changes greater than 1.00 in T cells and macrophages compared to all other generic cell types were retained. In total, 204 and 180 genes were removed prior to sub-clustering analysis of T cells and macrophages, respectively.

### Sub-clustering of Granuloma unified T and NK cells

Across the complete set of 44,766 T and NK cells, Louvain clustering was initially performed at a range of resolution of values (0.30 – 0.75) to examine the relationships between cluster membership. In this analysis, a cluster was observed to be defined by persistent expression of contaminating transcripts derived from macrophage and mast cells (Cluster 4 - Louvain Resolution 0.60). To confirm that these cells did not represent persistent doublets, all T cells were scored by expression of the top 20 cluster defining T cells and similar signature scores between the contaminated cell population were observed. Additional sub-clustering within the “contaminated” T cell cluster was performed to understand whether residual contamination obscured additional T cell biology; this failed to reveal additional T cell clusters not identified among the remaining non-contaminated populations. Since this contamination cluster was not observed to obscure a novel T cell phenotype, this population was excluded from downstream analysis. Following removal of the cluster of T cells defined by residual contamination, dimensionality reduction and clustering at multiple clustering resolutions (Louvain resolution: 0.25 – 0.75) were performed. In this final analysis, a total of 12 T cell populations were identified at a clustering resolution of 0.75. Finally, additional sub-clustering was performed within the population of 2,377 γδ and cytotoxic T cells, including dimensionality reduction and clustering at multiple resolutions (0.30 – 0.75). Here, 2 primary populations of cells were identified: sub-cluster 2, a population of cytotoxic cells enriched for expression of *TRDC* and sub-cluster 3, a population of XCL1+ NK cells. Differential expression analysis was performed to determine differences in gene expression between these clusters upon which the classification of these cells was based.

Additional sub-clustering analysis was performed within the T1-T17 population through repeated variable gene identification, dimensionality reduction and Louvain clustering (Resolution = 0.55), and 4 distinct sub-populations were discovered. Differential expression analysis was performed within the 9,234 T1-T17 cells using a Wilcox test in Seurat to identify sub-cluster defining gene signatures.

### Annotation of T /NK subclusters

T cell populations were classified using a combination of manual curation and comparison to literature-derived sequences. Granuloma T cell populations were compared to publicly available T cell population and scRNA-seq signatures. Specifically, comparisons were performed in the following ways:

1. For each T cell cluster, cluster-defining genes were compared to publicly available databases of immune signatures, including IPA, GeneGO, MSigDb (Liberzon et al., 2011) and SaVant (Lopez *et al*., 2017). This was performed by comparing the set of T cell cluster-defining genes (Adjusted p-value < 0.001 and log-FC > 0.2) to the signatures in GSEA and the SaVant data using Piano (Lopez *et al*., 2017; Varemo et al., 2013). Specifically, significance was assessed using a hypergeometic test to examine the likelihood of the observed frequency of enriched genes. Among cluster-defining genes for each T/NK cell sub-cluster, comparisons were performed within each GSEA collection C1-7 (https://www.gsea-msigdb.org/gsea/msigdb/collections.jsp) and to the SaVant database. Expression signatures were also compared to MSigDB signatures using GSEA. Here, pseudo-bulk expression signatures were generated for each T/NK sub-population as the average gene expression across all cells within each cluster. These average expression values were used to perform GSEA for each cluster in which the expression values were compared to all other clusters using 1,000 permutations.
2. Each T cell cluster was compared to literature-derived signatures of T cells from another scRNA-seq study. Here, cell signature scores were generated in Seurat using the AddModuleScore function using gene expression signatures obtained from human lung cancer (Guo et al., 2018). To determine the significance of these score, 1,000 permutations were performed in which T cell cluster identity was randomly re-assigned to generate a null distribution of module scores.
3. Finally, extensive manual curation was performed based on literature evidence. For each cell population, an extensive literature search was performed to support classification of T cell sub-populations based on patterns of enriched gene expression. For example, regulatory T cells were identified on the basis of expression of known regulatory T cell markers (*FOXP3, IKZF1*, and *TNFSF18/GITR*). However, in many cases, surface markers used to define canonical T cell populations were not detected in the scRNA-seq data.

Next, expression of *TRAC* and *TRBC* or *TRDC* was evaluated within T cells in the scRNA-seq data and the frequency of cells expressing either *TRAC/TRBC* (yellow) or *TRDC* (green) within each of the 13 clusters was calculated. While *TRAC/TRBC* expression was observed in all 13 subclusters, *TRDC* expression was observed mainly in subclusters 1-3 compared to subclusters 4-13. Finally, cluster-specific expression of *CD4* and *CD8A* and *CD8B* were examined as the proportion of cells with non-zero expression of *CD4*, *CD8A/B* or *CD4&CD8 (A/B)*.

### Sub-clustering of Granuloma Macrophages

Across 27,670 macrophages, dimensionality reduction and Louvain clustering at multiple clustering resolutions was performed. In initial clustering, a cluster defined by contaminating transcripts derived from other cell types (including mast cells (*KIT* and *CLU*), T cells (*CD3D*), and plasma cells (*JCHAIN*)) and soup-defining gene expression was identified. By comparing the distribution of macrophage-defining gene expression in this cluster to other clusters, this cluster was observed to have enriched signature scores relative other clusters. The enrichment of macrophage expression signatures was examined to determine the population of macrophages that have a core macrophage expression program. While this population of macrophages is primarily soup-defining gene expression, this cluster was not excluded due to the possibility that this represents an efferocytotic macrophage population.

### Classification of Macrophage Populations

Identities of the macrophage clusters were established through a combination of manual curation and comparison to published gene expression signatures from both population and scRNA-seq studies. More specifically:

1. For each macrophage cluster, similar comparison to databases of immune signatures including MSigDb and SaVanT were performed (See ***Identification of T cell Populations***).
2. A series of gene expression signatures were generated from published scRNA-Seq studies of macrophage states. For example, a recently published atlas of myeloid states in lung (Zilionis et al., 2019) was used to score granuloma macrophages. Further, a list of myeloid expression signatures was generated using lung myeloid cells from the Mouse Cell Atlas (Han *et al*., 2018). For each study, signatures for the top 20 cluster-defining genes were selected to generate gene expression signatures **(Table S5).** Signature scores were generated for each cell using the AddModuleScore function in Seurat.
3. Finally, in cases where an existing description of a macrophage population was not discovered, extensive literature searches were performed to contextualize possible identities of macrophage populations.

### Deconvolution of Bulk RNA-Sequencing Data

Population deconvolution was performed using CiberSort (Newman et al., 2015) using reference populations generated from random sampling of a quarter of the single cells within each of the 13 generic cell types identified in our single-cell analysis.

### Co-variation in Granuloma Composition

We calculated correlations in cell-type proportions to identify underlying structure in the co-occurrence of cell types across all granulomas. Specifically, we calculated Pearson correlation coefficients for all pair-wise cell-type combinations (**NB** we also performed each analysis using Spearman correlation coefficients and obtained similar results). For each pairwise combination of cell types, we calculated permutation p-values by randomly re-assigning cell type labels to generate a set of background correlation values (**Table S9,S10**)

We then performed hierarchical clustering to identify clusters of correlated cell-types across granulomas, calculating the proportional composition of correlated cell-type clusters within each lesion. For each of the 5 clusters identified through hierarchical clustering, we calculated permutation p-values to examine average correlation values. To understand the relationship between identified cell-type clusters and granuloma-level bacterial burden, we examined the abundance of correlated cell types by grouping lesions by timing of granuloma formation.

### Cell-Communication Analysis

To examine cell-cell interactions, we first generated a curated list of receptor-ligand pairs through a combination of publicly-available databases and literature review. Within each granuloma, we generated edge weights between cell types for a given receptor ligand pair by multiplying the average receptor expression in Cell Type 1 by the average ligand expression in Cell Type 2. Edge weights were constructed for all receptor-ligand pairs and pairwise-cell type combinations within granulomas individually. Within each granuloma, we performed a total of 1,000 permutations for each receptor-ligand pair in which cell-type identifiers were randomly resorted and the resulting edge weight was recorded. For each receptor-ligand pair, the significance of the observed value was calculated from a z-score comparison of the observed value relative the permuted values.

We further performed adjustment of receptor-ligand edge weights at multiple levels. (1) To account for differences in the relative abundance of ‘sender’ cell types, we multiplied receptor-ligand edge weights by the proportion of all ‘sender’ cells within a granuloma. In effect, this generates a pool of ‘sender’ cell derived ligand that is available to act upon cell types bearing appropriate receptors. (2) To identify the most likely receiver cells, we weighted receptor-ligand edge-weights by the proportion of total receptor expression within the receiving cell subset cluster relative to the average receptor expression across all cells in the granuloma. In this scheme, receptors with more uniform expression across the entire granuloma will be down-weighted to reflect non-autonomous sinks of extracellular ligands, while receptors predominantly expressed by a single cell subset will be up-weighted. (3) Finally, we adjusted receptor-ligand edge weights to account for the percent of cells within the receiver cell subset expressing a given receptor by multiplying our receptor-ligand edge weights by the proportion of all ‘receiver’ cells expressing the receptor within a the receiver cell subset.

To identify axes of intercellular communication with differential weights across granulomas, we performed student’s t-tests of receptor-ligand edge weights between (A) high-burden and low-burden lesions, and (B) original and late-blooming lesions. We filtered results based on the following criteria: (1) the average permutation p-values for the receptor-ligand pair within high or low-burden lesions < 0.05, (2) p-value from student’s t-test in (A) or (B) above < 0.05, and (3) fold-change of the adjusted receptor edge-weight > 0 in the (A) or (B) comparisons. The “dplyr” package in R was used to filter the resulting cell-cell interaction database to count significant interactions across cell type groups and granuloma burdens, identify cell type groups contributing to the top 10% of ligands most strengthened in either high or low burden granulomas, identify ligands most associated with high or low burden granulomas, and identify cell type specificity of these ligands. The “circlize” package in R was used to generate circus plots of the topology of signaling networks across high and low burden granulomas.

### Statistical methods

Non-parametric Spearman’s rho was calculated for correlation analysis for evaluating the degree of relationship between cellular abundance and bacterial burden. Non parametric t-test was used when comparing two groups (Mann-Whitney U). P values, or where appropriate adjusted or permutation p values, ≤ 0.05 were considered significant. Statistical analysis was performed using GraphPad Prism v8 (GraphPad software, San Diego, CA), JMP Pro v12 and R base statistics.

## Table legends

**Table 1:** T/NK subclusters characteristics and annotations

**Table S1:** Granuloma CFU, CEQ, CFU/CEQ; PET-CT: SUV-R, Size and Time of detection

**Table S2a:** Seq-Well array loading densities and doublet rate

**Table S2b:** Technical correction data: SoupX

**Table S2c:** Doublet removal Metadata

**Table S2d:** Cell level metadata

**Tablet S3:** Canonical cell type enrichment gene list: 13 cell type clusters

**Table S4:** Cell type composition: percentage of assigned granuloma cells. A) canonical cell type clusters, b)macrophage subclusters, c) T/NK subclusters and d) T1T17 subpopulation

**Table S5:** Correlation (Spearman’s rho) with bacterial burden and difference between in percentage of cells in early high burden and late low burden granulomas (Mann Whitney U): A) canonical cell type clusters, b) T/NK subclusters and C) T1T17 subpopulation

**Table S6:** T/NK subclustering: enrichment gene list :13 T/NK subclusters

**Table S7:** Type1-Type-17 subpopulation enrichment

**Tablet S8:** Macrophage subcluster enrichment: 9 subclusters

**Table S9:** Cellular ecology

**Table S10:** Cellular ecology correlation permutations (Spearman and Pearson)

**Table S11:** Association of cell group abundance with bacterial burden: (1) All: CFU low vs high, (2) timing of granuloma detection (Early vs late)

**Table S12:** Interaction analysis: receptor-ligand senders and receivers in early and late granulomas.

---

**Figure S1.**
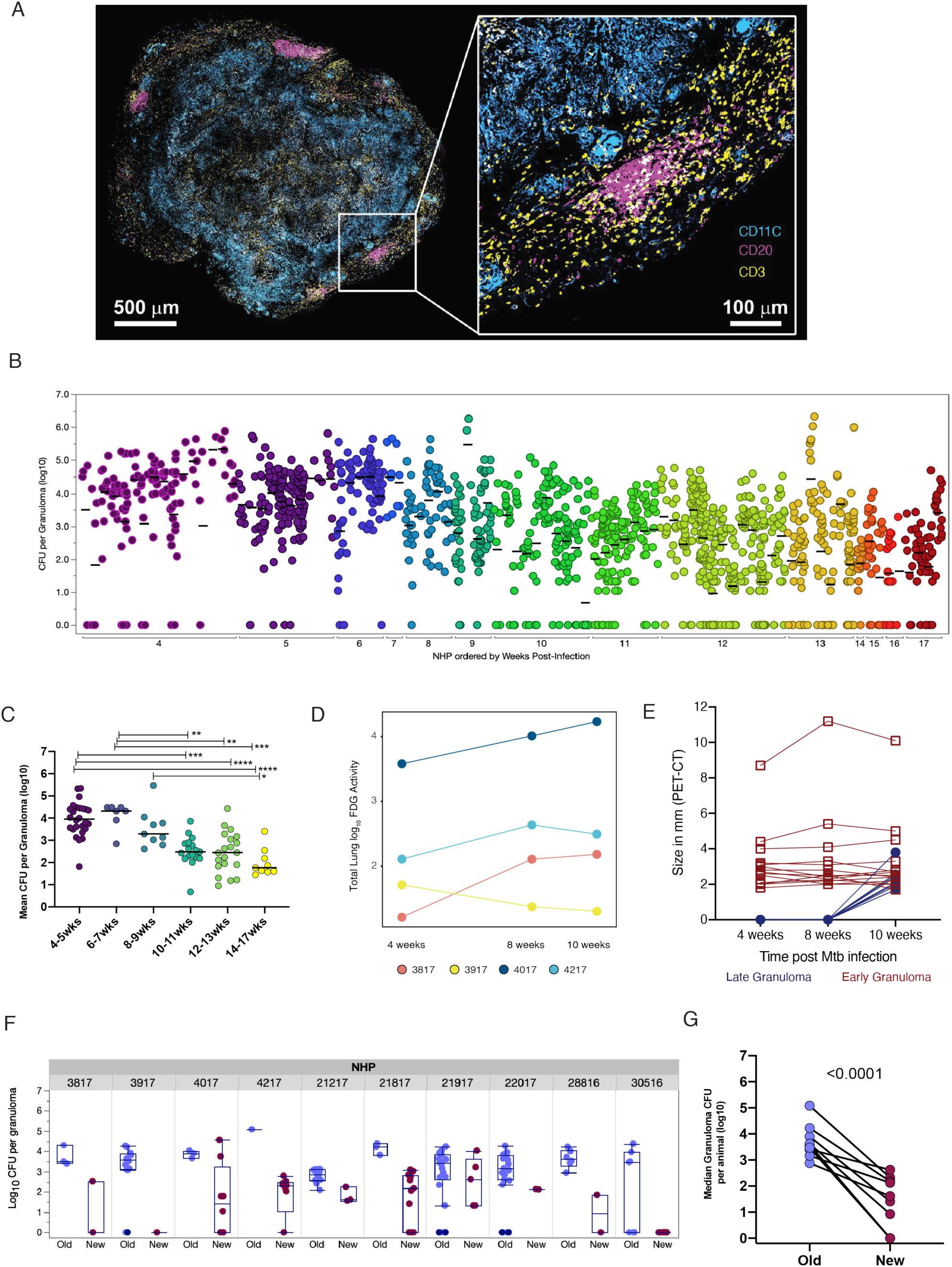
Granuloma architecture and CFU per granuloma decreases over time. (**A**) Architecture of macaque TB lung granuloma, where lymphocytes and macrophages are present in distinct regions. Immunohistochemistry and confocal microscopy were performed on a granuloma from an animal at 11 weeks post-Mtb infection to visualize localization of CD11c+ macrophages (cyan), CD3+ T cells (yellow), and CD20+ B cells (magenta). (B) Each column depicts the CFU for all granulomas of an individual macaque (N=88 macaques), ranging from 4 weeks to 17 weeks post-infection. Each dot represents a granuloma. Lines are at means (per animal) and different colors represent weeks post-infection. (C) CFU per granuloma decreases significantly starting at 10-11 weeks post-infection. Each dot represents the mean CFU per granuloma of an individual animal, with the x-axis indicating weeks post-infection at which necropsy was performed. Lines are at medians. Differences between time points were tested using Kruskal-Wallis test with Dunn’s multiple comparison adjustment. (* p < 0.05, ** p < 0.01, *** p < 0.001, **** p < 0.0001.) (D) Total lung FDG activity (in log scale) measured by PET scans of each animal at 4, 8 and 10-weeks post-Mtb infection showing trajectories of lung inflammation. (E) Size of each granuloma measured by CT scans at 4, 8 and 10 weeks post-mtb infection. Early granulomas are those identified at 4 weeks post infection (in maroon) and late granulomas are those identified at 10 weeks post infection (in dark blue). (F) CFU per granuloma is shown for early detection (blue) and late detection (red) within each animal. Box plots lines represent the median, IQR and range Each dot represents a granuloma. (G) CFU is significantly lower in new granulomas within animals. Each dot (and line) represents the median CFU per granuloma of each animal. Statistics: paired t-test

**Figure S2:**
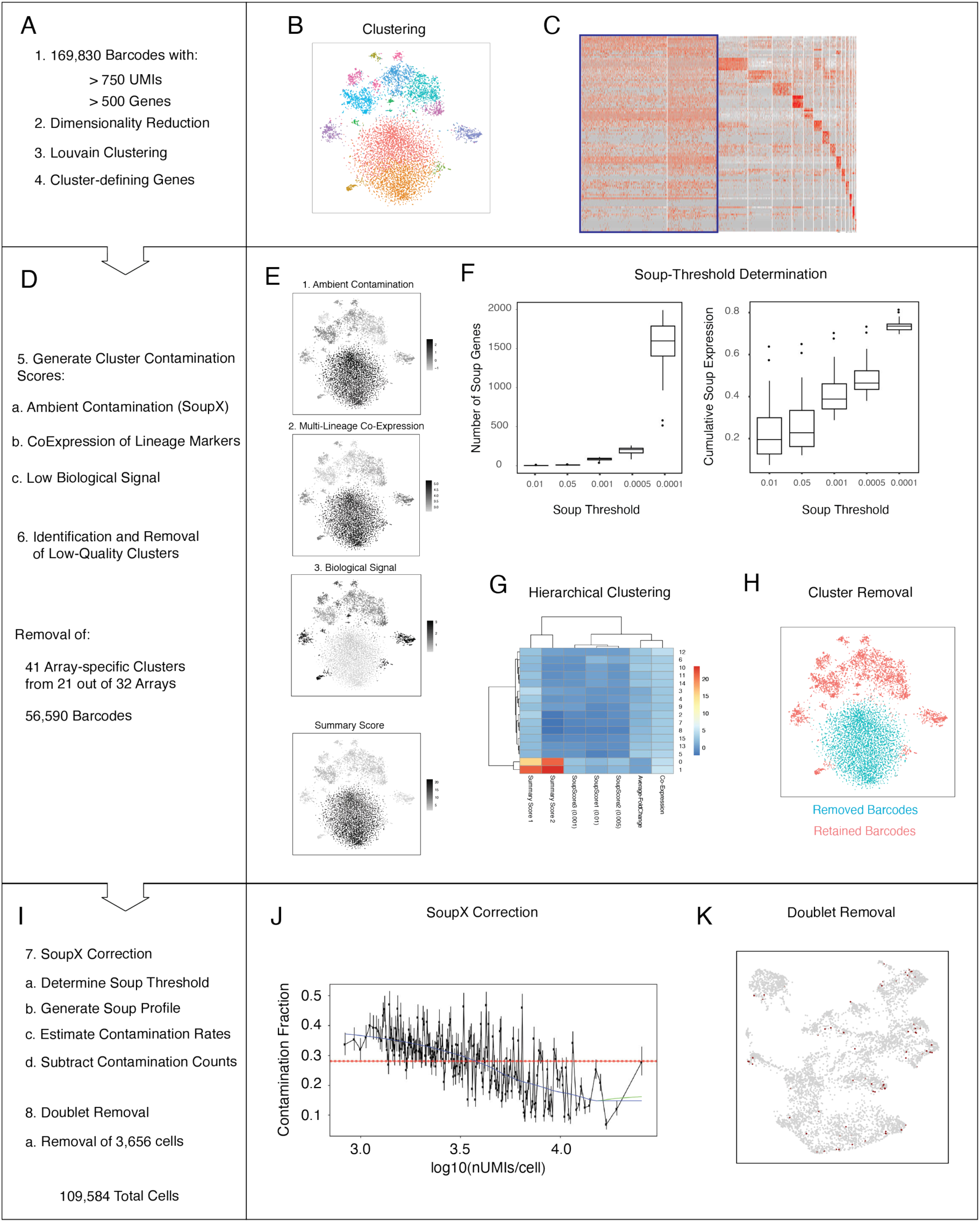
Sequencing, alignment and QC pipeline (see methods) **(A, D, I)** Array-specific processing pipeline. **(B)** Array specific Louvain clustering (Resolution = 1.25). **(C**) Cluster-defining gene expression was determined within each array. **(E)** Overview of Cluster-Specific Summary Score. **(F**) Estimation of soup-thresholds for correction of ambient RNA contamination. Left: Relationship between soup-thresholds (x-axis) the number of soup defining genes detected for each array (y-axis). Right: Relationship between soup-thresholds (x-axis) and the cumulative proportion of soup-defining gene expression (y-axis). **(G)** Hierarchical clustering results used to identify and remove clusters defined by ambient contamination from each array. **(H**) t-SNE plot showing removal of clusters characterized as ambient RNA. **(J)** Estimation of array-specific contamination rates using SoupX. **(K)** Identification and removal of array-specific doublets.

**Figure S3:**
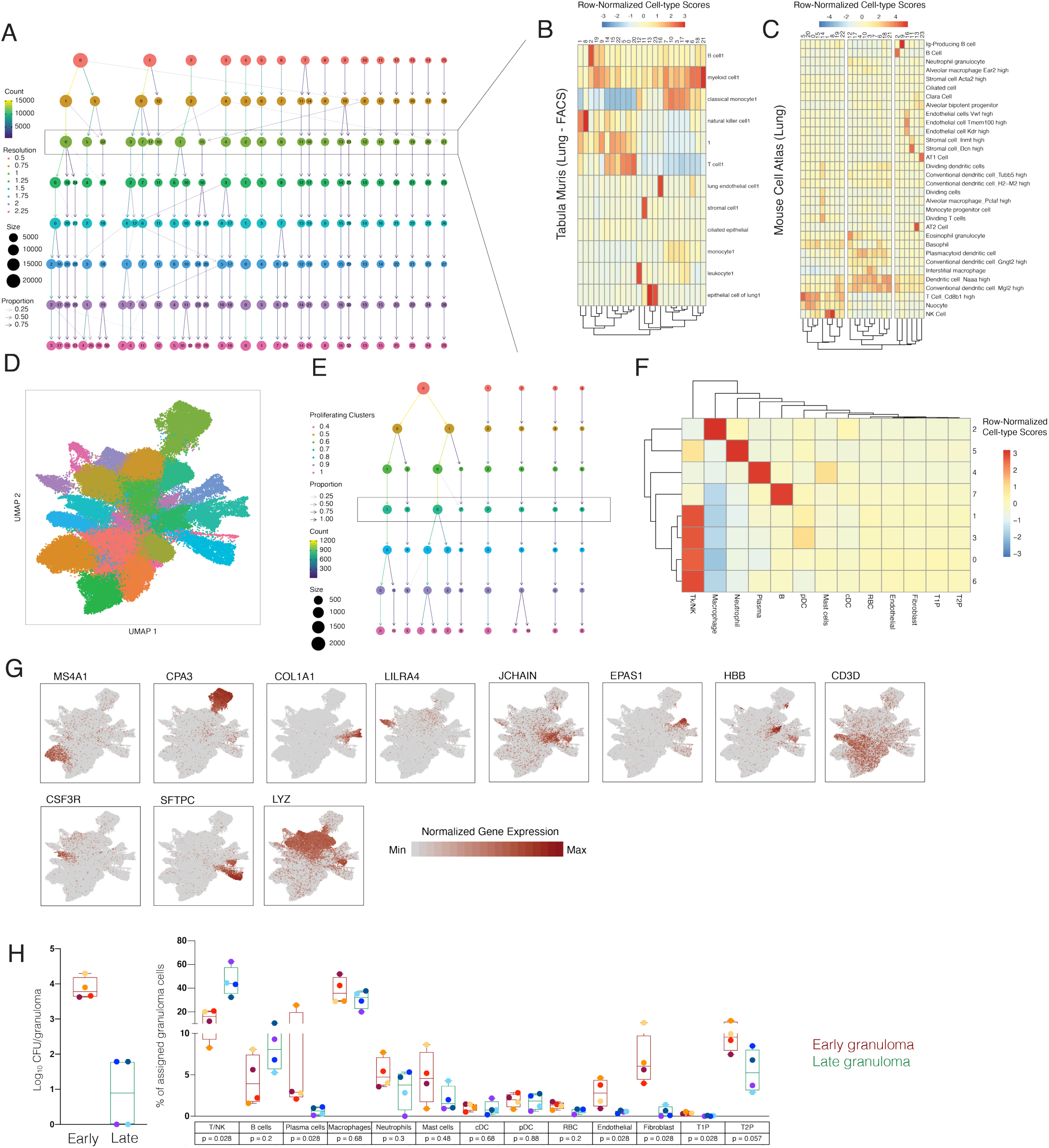
Identification of Canonical Cell Types. **(A)** Waterfall plot showing stability of cell-type clustersat multiple clustering resolutions. Boxed row (resolution=1.00) selected for downstream analysis. **(B, C)** Distribution of lung cell-type signatures obtained from the Tabula muris (B) and Mouse cell (C) atlas. **(D**) UMAP plot of 109,584 cells colored by Louvain clusters (resolution = 1.00). **(E)** Waterfall plot showing the stability of sub-clustering analysis of 3,123 cells with a proliferating gene signature. **(F)** Distribution of canonical cell type signatures across subclusters of proliferating cells. **(G)** Expression levels of cluster-defining genes overlaid on UMAP plot in panel 2A. (**H) Left:** CFU per granuloma based on the timing of detection by PET CT scan in one animal : 4017. **Right:** Difference in granuloma proportional composition of cell type clusters between early (maroon box plot) and late granulomas (green) within an animal (4017). Each granuloma is coloured. Statistics: Mann Whitney U. p values are presented in boxes. Box plot showing median, IQR and range; each dot represents a granuloma.

**Figure S4.**
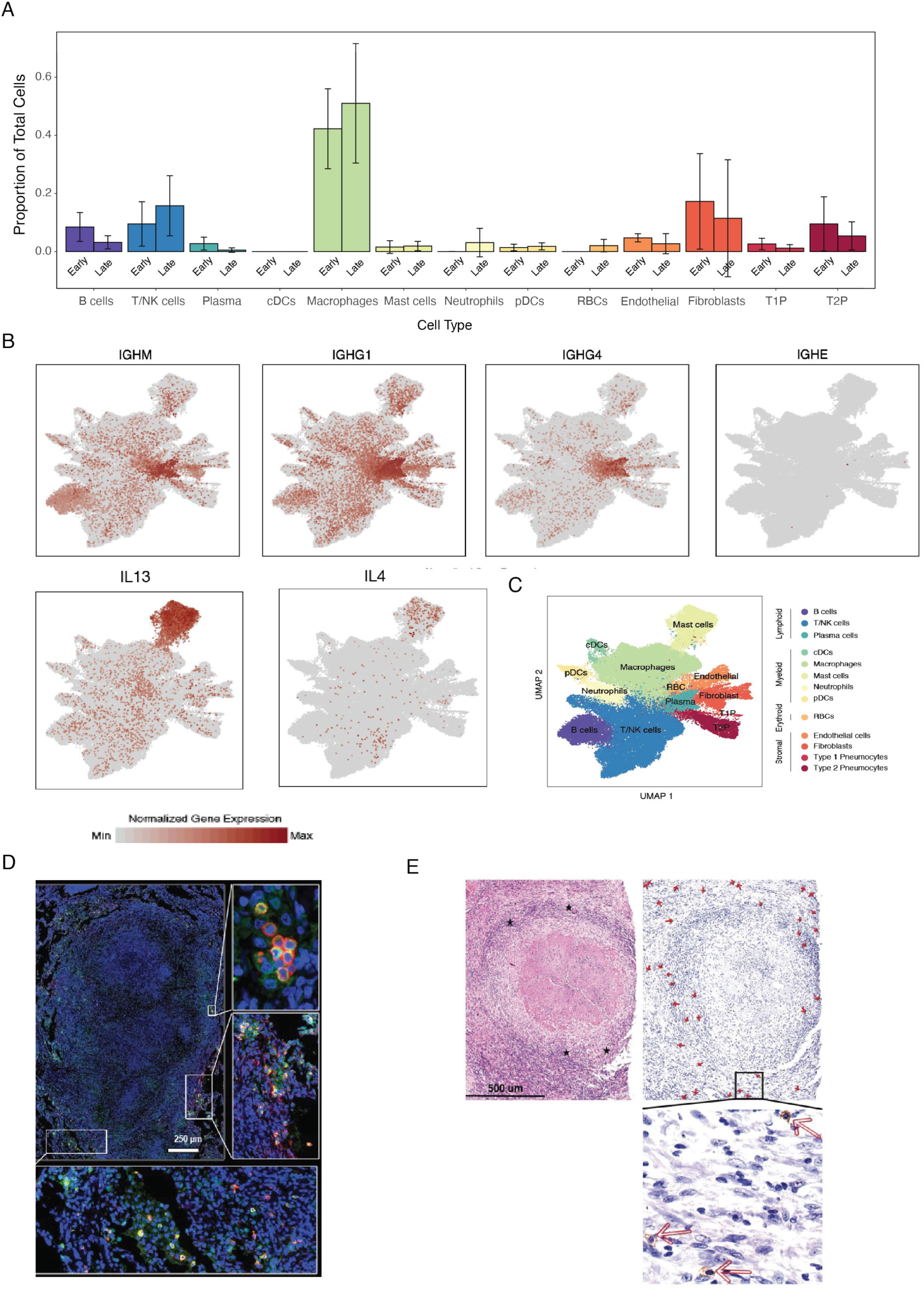
Cell type confirmation and Expression of selected functional transcripts. **(A)** Proportion of cell types in granulomas from bulk sequencing of 6 early and 6 late granulomas to confirm the trend seen in scRNAseq. **(B)** UMAP plot of 109,584 cells from 26 granulomas colored by identities of 13 generic cell types. **(C)** Expression levels of select functional genes overlaid on UMAP plot of 109,584 cells. **(D)** Detection of mast cells in a 10-week NHP granuloma using immunohistochemistry, staining for tryptase (green) and c-kit (CD117)(red). **(E)** Detection of mast cells in a human lung granuloma. Hematoxylin and eosin stain and immunohistochemistry with multinucleated giant cells (stars, (top left) and c-kit (CD117) staining (indicated by arrows, top and bottom right).

**Figure S5.**
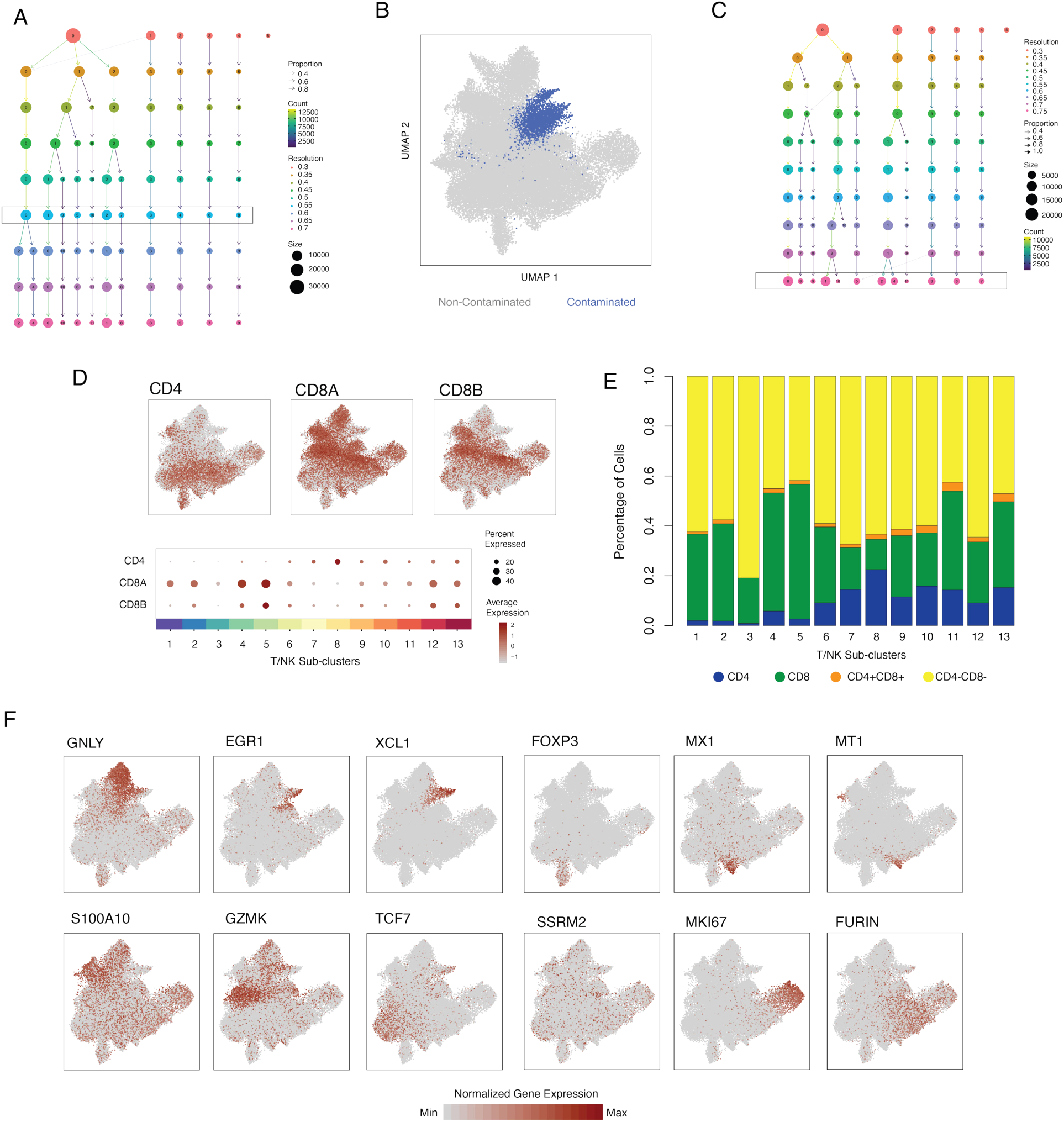
Sub-clustering and phenotypic identification of T/NK cell populations. (**A**) Waterfall plot showing the stability of T/NK cell sub-clustering. Boxed row (resolution=0.55) selected for downstream analysis. **(B)** UMAP plot of 44,766 T/NK cells with a sub-cluster of 3,544 T/NK cells defined by residual contamination highlighted (blue). **(C)** Waterfall plot showing the stability of T/NK cell sub-clustering following removal of contaminated T cell sub-cluster. Boxed row (resolution=0.75) selected for downstream analysis. **(D)** T/NK subclustering UMAP overlaid with normalized gene expression for CD4, CD8A, and CD8B (top). Expression of these genes across 13 sub-clusters (bottom) where color intensity corresponds to level of gene expression and size of dots represents the percent of cells with non-zero expression in each cluster. **(E)** Frequency of expression of *CD4* (blue), *CD8A* and/ *CD8B* (green), *CD4* and *CD8A/B* (orange) or no expression of *CD4/CD8A/B* (yellow) across 13 T/NK cell subclusters. **(F)** UMAP plots overlaid with normalized expression levels for selected T/NK cell subcluster-defining genes.

**Figure S6:**
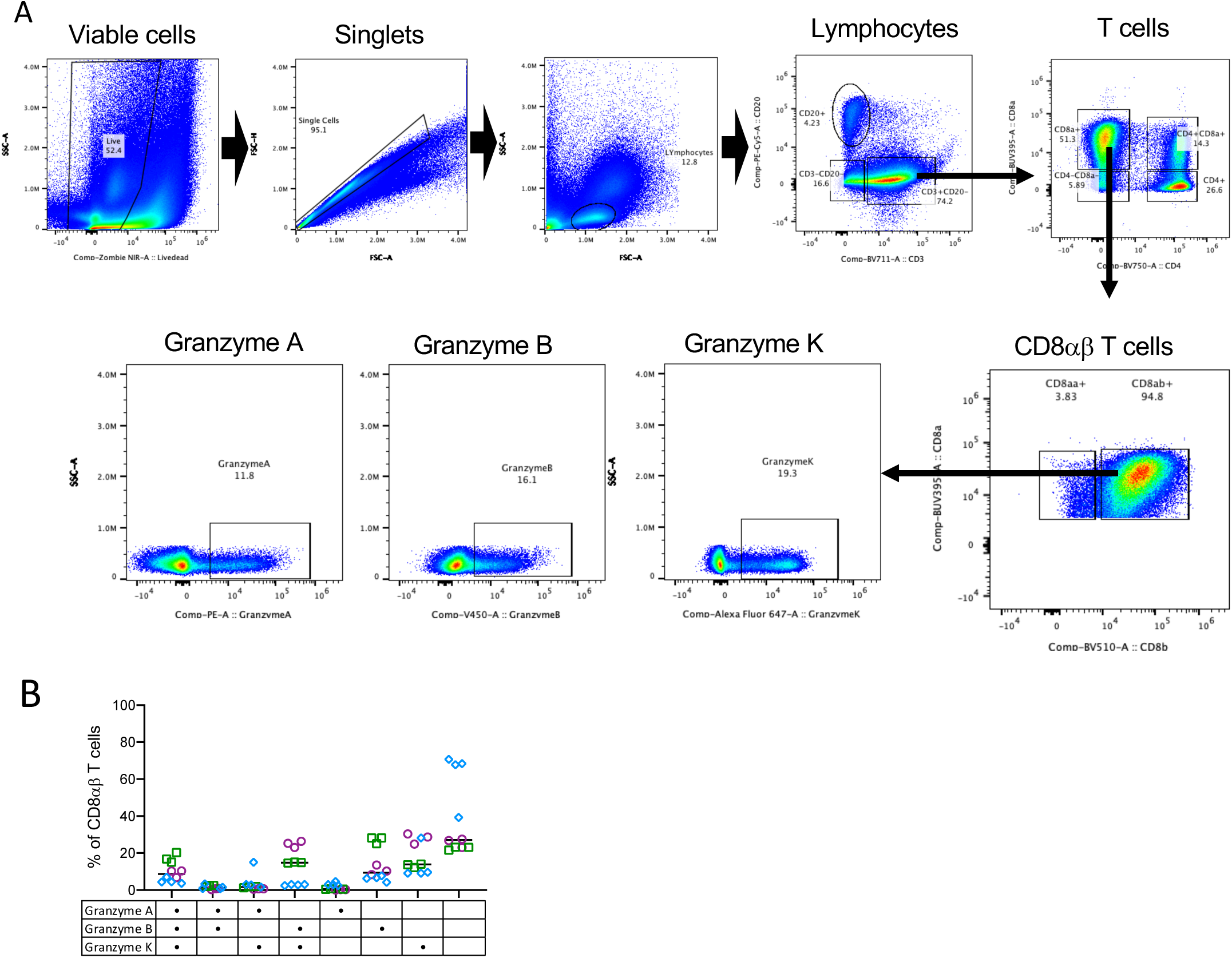
Flow cytometry confirmation of cytotoxic molecules in TB granulomas. (**A**) Gating Tree showing identification CD8ab T cells in lung granuloma samples and population of Granzyme A, Granzyme B and Granzyme K + CD8abT cells. **(B)** Frequency of CD8ab T cells in lung granulomas making one or more (two , three) types of Granzymes (A, B or K). Each symbol is a granuloma and each colour identifies an animal. This data supports different types of granzyme producing cytotoxic cells identified in scRNAseq.

**Figure S7.**
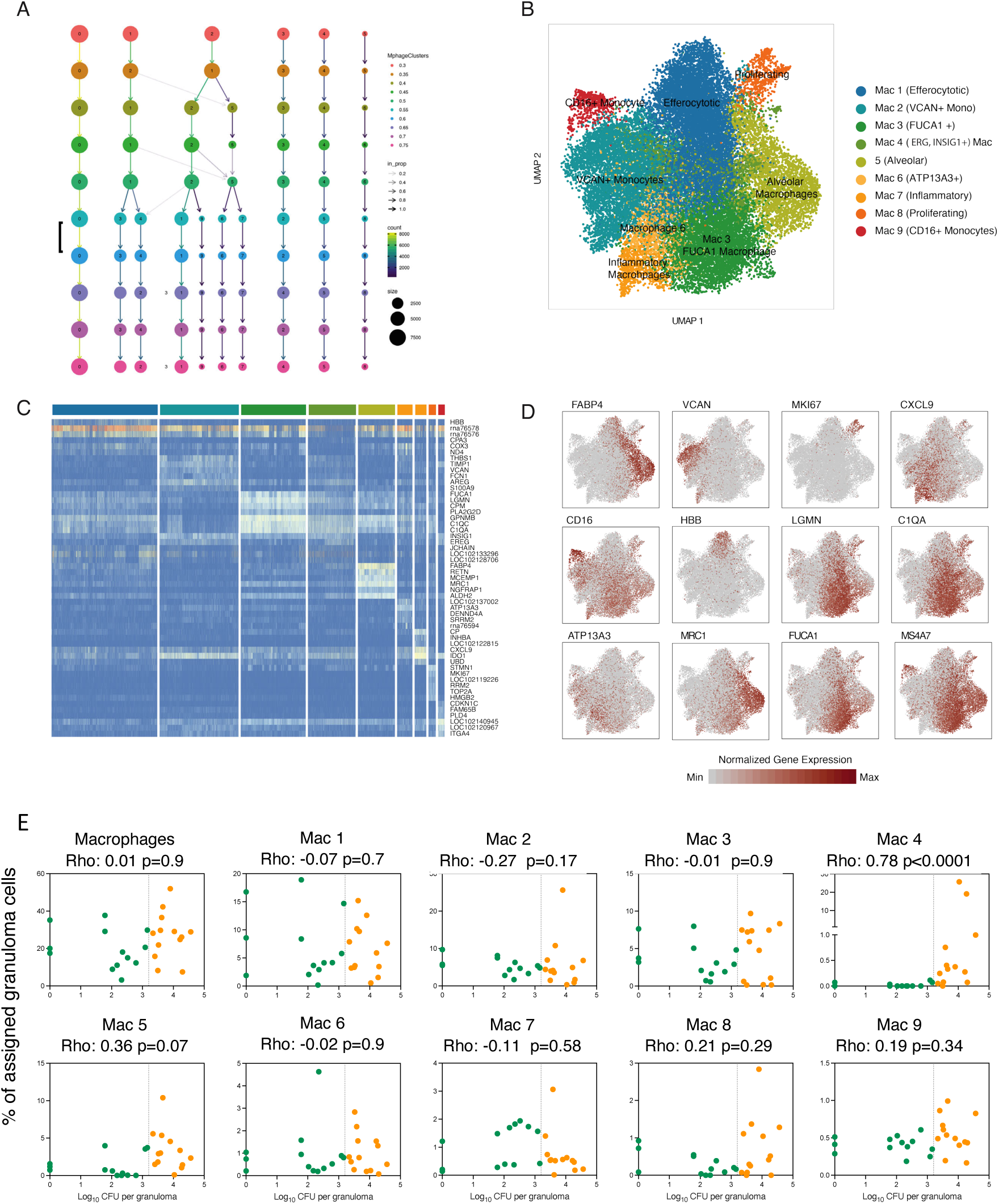
Macrophage heterogeneity in Mtb granulomas. **(A)** Waterfall plot showing the stability of macrophage sub-clusters. Boxed row (resolution=0.55) selected for downstream analysis. **(B)** UMAP plot of 27,670 macrophage cluster colored by phenotypes. **(C)** Cluster-defining genes across macrophage subclusters. **(D)** Macrophage subcluster-defining genes overlaid on macrophage plot in panel B. **(E)** Significant correlations between proportion of Macrophage subclusters with bacterial burden of individual granulomas (Log10 CFU per granuloma) using non-parametric Spearman’s rho correlation test. Color indicated binned granuloma bacterial burden: low (green) and high (orange).

**Figure S8.**
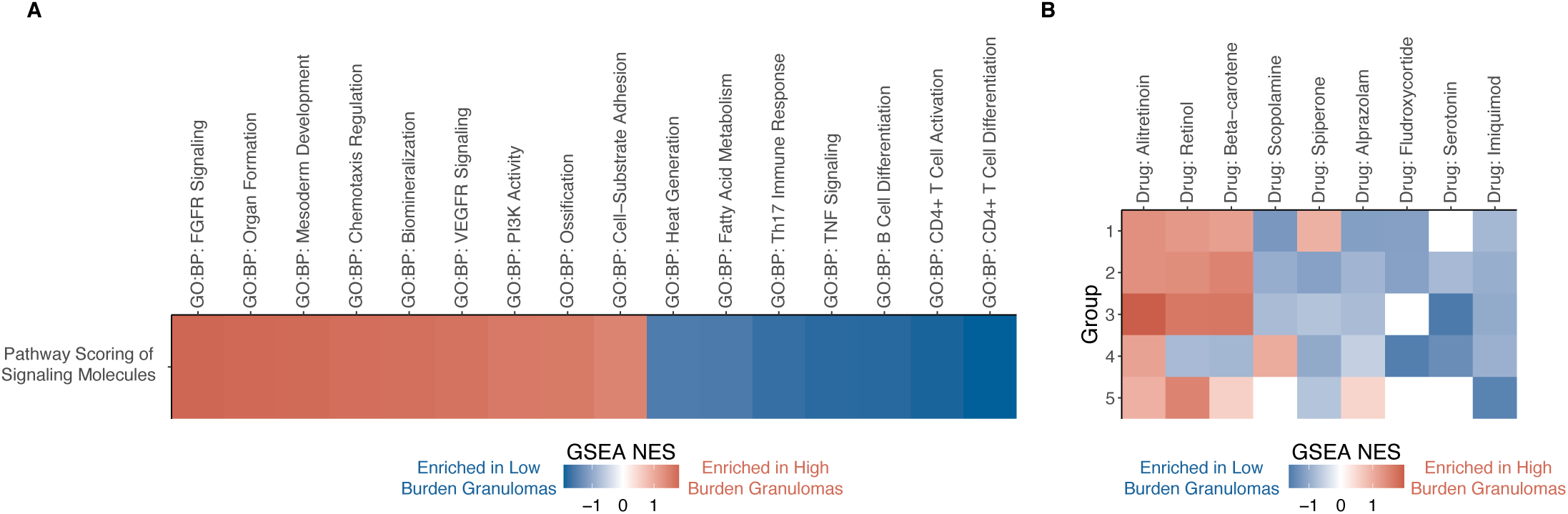
Transcriptomic pathways associated with granuloma burden. **(A)** Pathways enriched in signaling molecules associated with high vs. low granuloma burden. Signaling molecules were ranked according to their log(fold-change in high vs. low burden granulomas) as input to GSEA. **(B)** Drugs with targets enriched in signaling molecules associated with each cell type group.

